# Melatonin nanoparticles inhibit mutant hematopoiesis & restore bone marrow architecture in myeloproliferative neoplasms

**DOI:** 10.64898/2026.08.28.746520

**Authors:** Siddharth Gupta, Alessandro Motta, Sara Elsafy, Shiva Khorshid, Alessia Nucci, Vishrutha Sampath, Ananya Bhattacharjee, Margherita Vieri, Kathrin Olschok, Kristina Pannen, Jelena Lazarevic, Maria Jimena Rodriguez, Patrick Weiand, Venkatakrishnan Hariharan, Cristina Baquero López, Chunxiao Zhou, Henrike Jacobi, Bärbel Junge, Tata Nageswara Rao, Fabian Kiessling, Emiel P. C. van der Vorst, Twan Lammers, Federica De Lorenzi, Julian Baumeister, Steffen Koschmieder, Marcelo Augusto Szymanski de Toledo, Alexandros Marios Sofias, Nicolas Chatain

## Abstract

Myeloproliferative neoplasms (MPN) are chronic hematologic malignancies characterized by clonal myeloid expansion, inflammation, oxidative stress, and progressive bone marrow (BM) remodeling that may culminate in fibrosis and secondary acute leukemia. Here, we evaluated the therapeutic efficacy and the underlying mechanisms of melatonin (MT) and liposomal melatonin (nano-MT) in preclinical MPN models. MT selectively inhibited clonogenic growth of patient-derived peripheral blood mononuclear cells and induced pluripotent stem cell-derived CD34⁺ hematopoietic stem and progenitor cells in comparison to healthy controls. This effect was associated with increased apoptosis, reduced reactive oxygen species (ROS), and decreased glucose uptake, independently of MT receptor signaling. Transcriptomic profiling of primary MPN CD34⁺ cells revealed suppression of MYC targets, G_2_M checkpoint signaling, ROS, and glycolysis pathways. In co-culture models, MT reduced stromal α-smooth muscle actin and phosphorylated SMAD2/3, indicating inhibition of TGF-β-driven mesenchymal stromal cell-to-myofibroblast formation. In tamoxifen-inducible SclCreER;JAK2^V617F^ mice, nano-MT achieved efficient spleen and BM targeting. Therapeutically, nano-MT reduced erythrocytosis, myeloid progenitor expansion, and BM IL-1β levels. Longitudinal micro-computed tomography and histological analyses demonstrated normalization of BM architecture, reduced osteosclerotic remodeling and splenomegaly, decreased reticulin deposition and megakaryocyte numbers. In a dose-escalation study, nano-MT restored erythrocyte, hematocrit, and platelet counts and normalized megakaryocyte-erythroid progenitors. Combination treatment with ruxolitinib further reduced leukocytosis, neutrophilia, and monocytosis. Collectively, these findings demonstrate that (nano-)MT attenuates MPN and BM remodeling by targeting metabolic, inflammatory, and fibrotic pathways. This study provides the first evidence for a therapeutic benefit of nano-MT in MPN and establishes a rationale for further translational evaluation.

**Key points:**

- Melatonin impairs MPN progenitors and blunts TGF-β-driven stromal activation *in vitro*.
- Liposomal melatonin normalizes blood counts, megakaryocyte burden, and bone marrow fibrosis in MPN mice.

## Introduction

Philadelphia chromosome-negative myeloproliferative neoplasms (MPN), comprising polycythemia vera (PV), essential thrombocythemia (ET), and primary myelofibrosis (PMF), are clonal disorders arising from mutant hematopoietic stem cells (HSC). Most cases are driven by a gain-of-function mutation in *Janus kinase 2* (*JAK2*) gene, followed, less frequently, by mutations in *Calreticulin* (*CALR*) and/or *MPL*, each of which drives expansion of myeloid progenitors.

Advanced MPN, and myelofibrosis in particular, are characterized by sustained inflammatory signaling, oxidative stress^1^ and pathological remodeling of the bone marrow (BM) stroma with deposition of fibrous extracellular matrix (ECM)^2,3^. These changes progressively impair normal hematopoiesis and establish a self-reinforcing niche that sustains the malignant clone^4–6^. Clonal megakaryocytes (MK) play a central role in this process exhibiting aberrant cytokine secretion profiles, including fibrogenic factors such as transforming growth factor-β (TGF-β), interleukin 1 beta (IL-1β), and platelet-derived growth factor (PDGF)^7–9^. Disease biology is therefore not confined to the mutant clone but is co-determined by the stroma it corrupts.

Current MPN therapies such as JAK1/2 inhibitors (e.g., ruxolitinib) and interferon alpha (IFNα) effectively control symptoms but are largely palliative, with limited impact on molecular remission or fibrosis reversal^10–12^. These limitations underscore the need for therapies that target both malignant HSC and the diseased microenvironment^13–15^.

Melatonin (N-acetyl-5-methoxytryptamine, MT) is a neurohormone predominantly produced by the pineal gland, classically known for regulating circadian rhythms^16^. However, it also exerts antioxidant effects by scavenging reactive oxygen species (ROS), anti-inflammatory effects by inhibiting NF-κB signaling, and antifibrotic effects in models of hepatic and pulmonary fibrosis by suppressing TGF-β mediated fibroblast activation^17–20^. Recent evidence further underscores the multifunctional role of MT as a modulator of bone homeostasis, mitigating osteoporosis, and structural degeneration but also safeguarding mitochondrial function and cellular ion balance^21^.

MT has been administered at high doses in both preclinical and clinical studies and is generally well tolerated even at supraphysiological levels, but it shows limited clinical efficacy due to unfavorable pharmacokinetic properties^16,22^. Accordingly, liposomal melatonin (nano-MT) has recently emerged as a pharmacologically optimized delivery formulation that can improve systemic exposure and tissue penetration^23–25^.

In this study, we evaluated the potency and selectivity of MT in primary and induced pluripotent stem cell (iPSC)-derived hematopoietic stem and progenitor cells (HSPC) from MPN patients and healthy donors (HD) in mono- and myeloid-stromal co-cultures. Furthermore, we tested free MT against ruxolitinib and liposome-encapsulated nano-MT *in vivo* in an inducible SclCreER;JAK2^V617F^ mouse model^26,27^, and employed a longitudinal multimodal imaging approach to verify the induction of the disease phenotype, visualize and quantify nano-formulation biodistribution, and assess therapeutic response.

## Methods

### Patient material

All MPN samples were collected at the Department of Hematology, Oncology, Hemostaseology and Stem Cell Transplantation at RWTH Aachen University Medical Center after patients’ written informed consent, as approved by the local ethics committee (EK 127/12). Both primary mesenchymal stromal cells (MSC), isolated from femoral heads^28^, and HD blood samples were provided by the Transfusion Medicine in the University Hospital, RWTH Aachen (EK 099/14, EK 300/13).

### Generation of iPSC-derived HSPC

iPSC derived from CALR-mutated MPN patient (iPSC line del52 het01) or healthy donors (HD, iPSC lines HDT4 and HD06) were cultured and subjected to spin-EB assays for HSPC generation as described previously^29^. On day 10-14, enrichment for CD34^+^ HSPC was performed by MACS following manufacturer’s protocol (CD34 MicroBead Kit, Miltenyi Biotec, Bergisch Gladbach, Germany) and used for downstream analysis.

### Animal Experiments

All animal experiments were planned and reported in accordance with the ARRIVE guidelines and were performed in compliance with the German Animal Welfare Act (TierSchG), the German Laboratory Animal Regulation (TierSchVersV), and EU Directive 2010/63/EU. The animal experiments were approved by the local authorities of North Rhine-Westphalia, Germany (LAVE). The sample size for animal experiments was determined using G*Power for a two-tailed unpaired t-test with α=0.05 and β=0.20 (80% power). All preparations and procedures are described in the Supplemental Methods, available online.

### Nanoformulation synthesis

Nanoparticles were prepared via microfluidics according to established procedures^23^. Details about synthesis and characterization can be found in the supplemental methods.

### Statistical analysis

All experiments were performed at least three times unless otherwise stated. Statistical differences were calculated using the tests as indicated in the figure legends, with P < 0.05 considered significant, and graphs were plotted using GraphPad Prism 11 (GraphPad Prism, USA).

## Results

### Melatonin selectively impairs clonogenic growth and induces apoptosis in MPN progenitor cells

To determine whether MT affects malignant hematopoiesis, we assessed its effects on clonogenic growth as a functional readout for progenitor proliferation and differentiation. In colony-forming unit (CFU) assays of peripheral blood mononuclear cells (PBMC) from HD and MPN patients, MT reduced colony numbers dose-dependently in MPN samples (**Fig. 1A**, **Fig. S1A**, **Table 1**). At 500 µM MT, colony numbers were significantly reduced in MPN compared with HD, indicating a selective sensitivity to malignant progenitors. This inhibitory effect was observed independent of genotype-specific differences. At 750 µM, both HD and MPN PBMC showed substantial reductions; therefore, 500 µM MT was selected for follow-up *in vitro* experiments. CFU-derived viable cell counts recapitulated the clonogenic growth reduction (**Fig. 1B**). To validate these observations in a genetically defined clonal model and reduce variability associated with patient-derived samples, we applied CD34⁺ HSPC enriched from HD- and MPN-derived iPSC to CFU assays using a single pulse of MT, alone or combined with IFNα. Treatment with MT alone or in combination with IFNα displayed minimal effect on colony numbers, whereas single IFNα treatment reduced colony numbers from HD (**Fig. S1B**). In contrast, MPN CD34⁺ HSPC exhibited a significant reduction in colony formation following MT treatment, comparable to IFNα (**Fig. S1B**). Total cell yields recovered from colonies treated with MT were largely unaffected in HD, while MPN-derived viable cells were markedly reduced by both MT and IFNα, as well as their combination (**Fig. S1C**).

**Figure 1.**
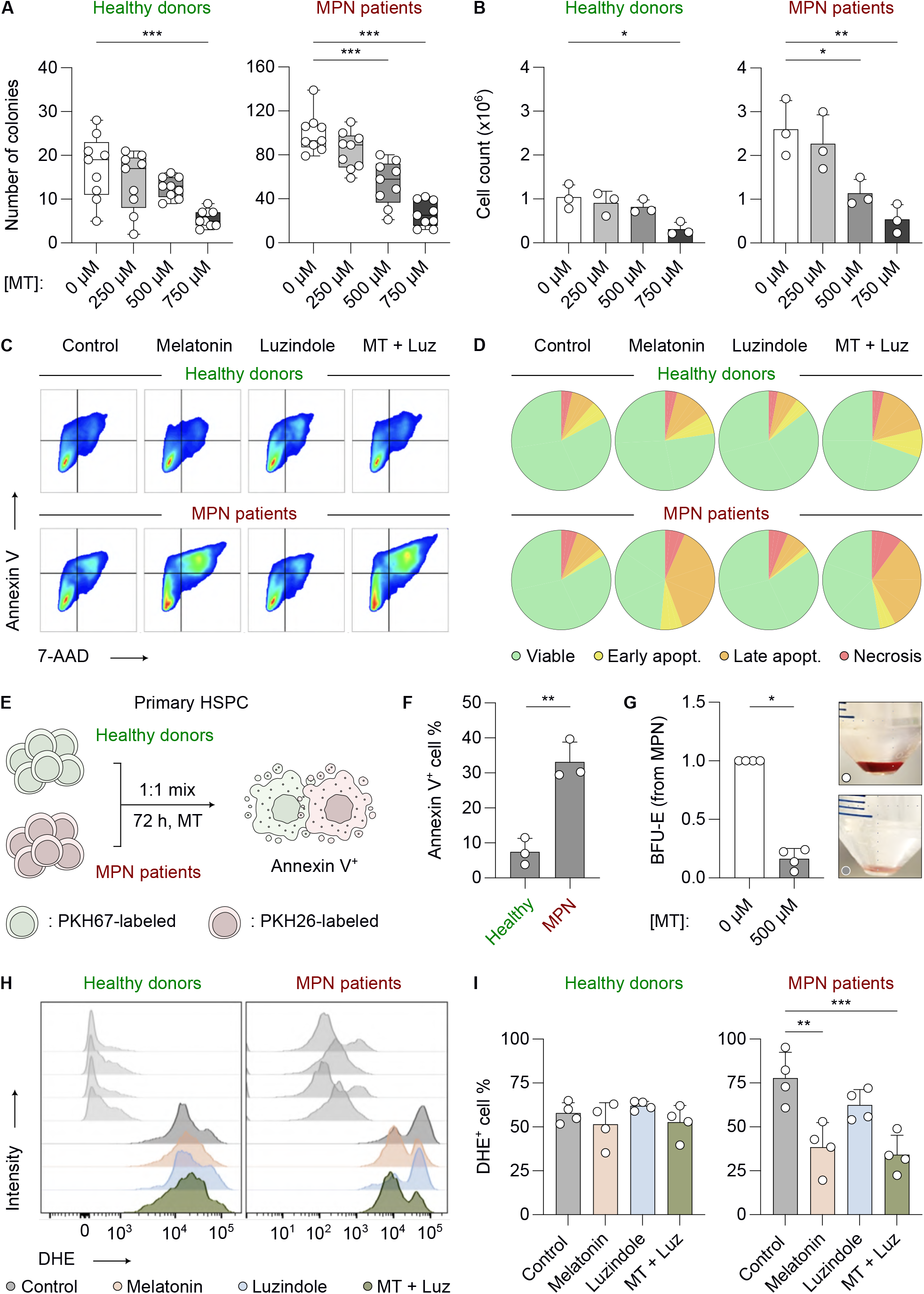
Melatonin selectively suppresses clonogenic growth, induces apoptosis, and reduces oxidative stress in MPN progenitors. **(A)** Total colony numbers in colony-forming unit (CFU) assays using PBMC from healthy donors or MPN patients treated with increasing concentrations of melatonin (MT; 0, 250, 500, or 750 μM) for 14 d. Box-whisker min-max plots show the median and interquartile range; n=3 biological replicates, n=3 technical replicates each. Statistical analysis was performed via ordinary One-way ANOVA with Dunnett correction for multiple comparisons against the control group. **(B)** Total viable cell counts recovered from the CFU assays shown in A. Bars show mean ± SD; n=3 biological replicates. Statistical analysis was performed via ordinary One-way ANOVA with Dunnett correction for multiple comparisons against the control group. **(C)** Flow cytometry contour plots of Annexin V versus 7-AAD staining in healthy donor- or MPN-derived iPSC-derived hematopoietic stem and progenitor cells (HSPC) treated with vehicle, MT (500 μM), luzindole (10 μM; MT receptor antagonist), or a combination of MT + luzindole for 72 h. **(D)** Quantification of viable, early apoptotic, late apoptotic, and necrotic populations from the experiment shown in C. Pie charts show the mean proportion of each population; n=3 biological replicates. **(E)** Schematic representation of the competitive co-culture apoptosis assay. Healthy donor-derived HSPC were labeled with PKH67, while MPN patient-derived HSPC were labeled with PKH26. The two populations were mixed at a 1:1 ratio before treatment with MT for 72 h and subsequent Annexin V analysis. **(F)** Percentage of Annexin V+ cells within the healthy donor- or MPN patient-derived HSPC populations following MT treatment as shown in E. Bars show mean ± SD; n=3 biological replicates. Statistical analysis was performed via an unpaired two-tailed t-test with Welch correction. **(G)** Burst-forming unit-erythroid (BFU-E) output from 4 independent PV samples following a single pulse of MT (500 μM). Data are expressed as fold-change relative to the corresponding control group. Bars show mean ± SD; n = 4 biological replicates. Statistical analysis was performed via a two-tailed one-sample t-test against the fold-change value of 1 for the control group. **(H)** Flow cytometry histograms, showing dihydroethidium (DHE; 25 μM) fluorescence in healthy donor- and MPN patient-derived iPSC-HSPC treated with vehicle, MT, luzindole, or combination of MT + luzindole. **(I)** Quantification of DHE+ cells under the indicated treatment conditions presented in H. Bars show mean ± SD; n = 4 biological replicates. Statistical analysis was performed via ordinary One-way ANOVA with Dunnett correction for multiple comparisons against the control group. Statistical significance is indicated by * (P < 0.05), ** (P < 0.01), *** (P < 0.001), and **** (P < 0.0001).

To investigate whether the reduction in clonogenic growth was associated with apoptosis, we performed Annexin V/7-AAD staining in HD- and MPN-derived CD34⁺ cells. In contrast to HD cells, MPN cells displayed an increase in Annexin V positivity upon MT treatment (**Fig. 1C,D**). Notably, co-treatment with the MT receptor antagonist luzindole (Luz) did not reverse this effect, indicating that MT induced apoptosis in a canonical receptor signaling-independent manner.

To evaluate the differential effects of MT on normal versus malignant hematopoietic progenitors within the same culture conditions, we established competitive co-culture assays in which HD-derived HSPC were labelled with PKH67 and PV-derived HSPC with PKH26 (**Fig. 1E**). Following 72 h of MT exposure, flow cytometric analysis demonstrated a significant increase in Annexin V⁺ apoptotic cells in the MPN cohort, whereas HD progenitors remained largely unaffected (**Fig. 1F**). Consistent with these observations, functional assays revealed a pronounced reduction in erythroid output, as evidenced by the loss of BFU-E colonies from PV patient-derived HSPC upon MT treatment (**Fig. 1G**). These results indicate that MT preferentially targets malignant progenitors by inducing apoptosis and suppressing erythroid colony formation, while sparing healthy HSPC.

Given the established role of oxidative stress in regulating HSPC fate^1^, we also examined whether MT affected intracellular ROS levels. DHE staining demonstrated no changes in HD progenitors across treatment conditions (**Fig. 1H-I**). In contrast, MPN-derived HSPC displayed elevated basal ROS, which were significantly reduced upon MT exposure (**Fig. 1H-I**). Neither Luz alone nor the combination of MT + Luz reversed this effect, further supporting a receptor-independent mechanism. MT thus preferentially targets MPN progenitors, and this selectivity is accompanied by normalization of the elevated redox state.

### Melatonin suppresses proliferative and metabolic pathways in MPN progenitors

To gain mechanistic insight into how MT affects malignant progenitors, we performed 3’-mRNA sequencing of MPN PBMC-derived CD34⁺ cells (**Fig. 2A**). Differential gene expression (DGE) analysis revealed 47 significantly downregulated and 15 significantly upregulated genes upon MT treatment (**Fig. 2B**). Pathway-level interrogation by Reactome analysis highlighted significant enrichment for processes involved in mRNA processing, mitotic progression (G_2_/M, S-phase), and organelle biogenesis among the MT-downregulated transcripts (**Fig. 2C**, **Fig. S2A**). Similarly, Gene Set Enrichment Analysis (GSEA) demonstrated suppression of proliferative signatures, including MYC, E2F, and G_2_M checkpoint gene sets, as well as reduced glycolysis and ROS-related pathways in MT-treated MPN cells (**Fig. 2D**). KEGG pathway enrichment analysis identified DNA replication, Glycolysis/Gluconeogenesis, and HIF-1 signaling among pathways enriched in MT-downregulated genes independent of MT receptor (*MTNR1A*, *MTNR1B*) regulation (**Fig. S2B,C**).

**Figure 2.**
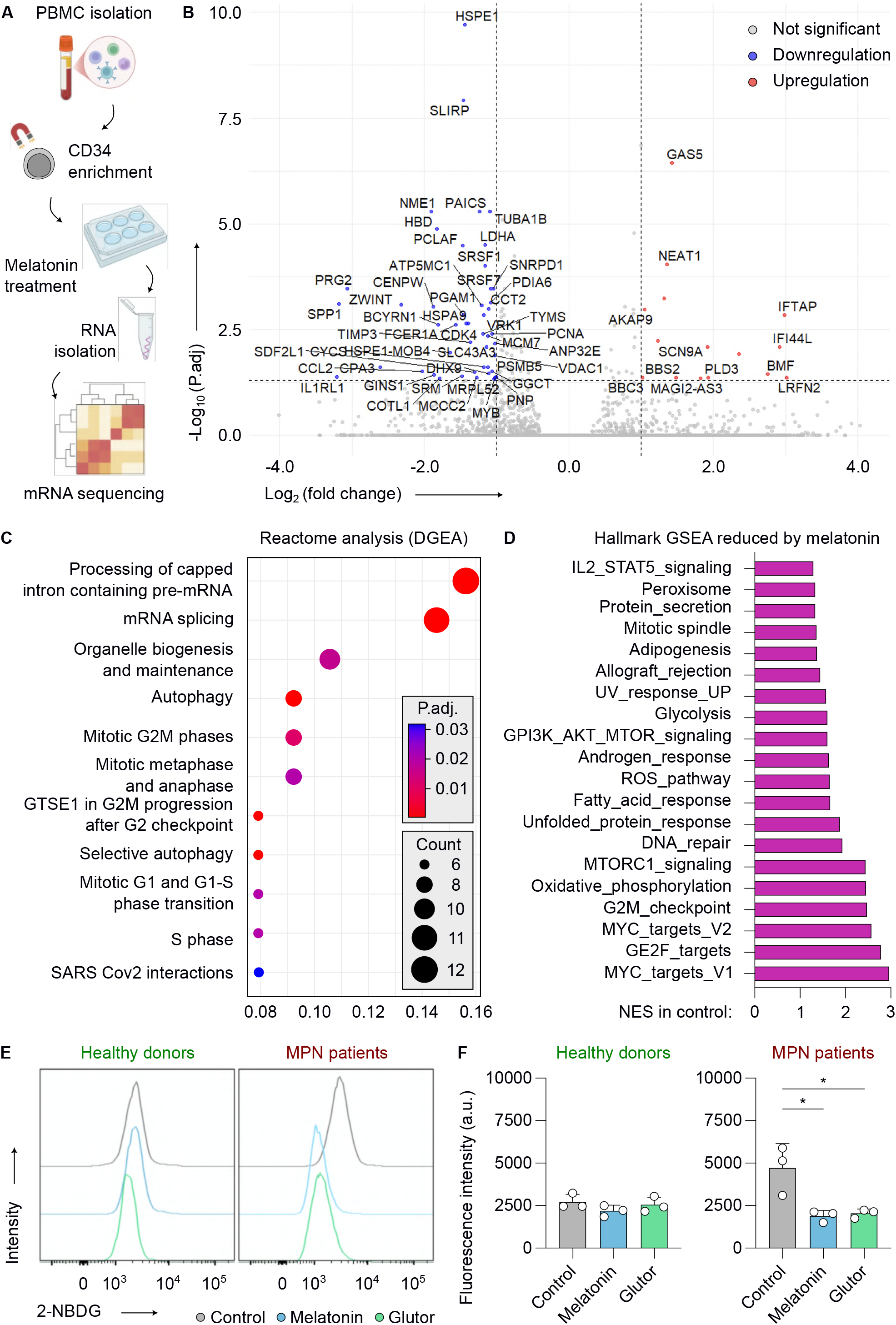
Melatonin suppresses proliferative and glucose-dependent metabolic programs in MPN hematopoietic progenitors. **(A)** Schematic representation of the experimental workflow. CD34⁺ HSPC derived from healthy donors (n = 3) or MPN patients (n = 6) were treated with vehicle or MT (500 µM) for 16 h, followed by RNA isolation and 3′ mRNA sequencing. **(B)** Volcano plot showing differential gene expression in MPN patient-derived CD34⁺ HSPC treated with MT compared with the corresponding vehicle-treated cells. Each point represents one gene. Genes with an adjusted P value < 0.05 and an absolute log_2_ fold-change > 1.5 were considered differentially expressed and are indicated as upregulated or downregulated. Differential expression analysis was performed using DESeq2 with Benjamini-Hochberg correction for multiple testing. **(C)** Reactome pathway-enrichment analysis of differentially expressed genes identified in panel B. Dot position indicates the gene ratio, dot size represents the number of genes contributing to each pathway, and color indicates the adjusted P-value. Pathway-enrichment P-values were corrected for multiple testing using the Benjamini-Hochberg method. **(D)** Gene set enrichment analysis of Hallmark gene sets in MT-treated versus vehicle-treated MPN HSPC. Bars show normalized enrichment scores (NES) for gene sets reduced following MT treatment. Gene sets with a false discovery rate q value < 0.05 were considered significantly enriched. Significance was assessed using 1000 gene-set permutations. **(E)** Representative flow cytometry histograms, showing uptake of the fluorescent glucose analogue 2-NBDG in healthy donor- and CALRdel52het01 MPN iPSC-derived HSPC treated with vehicle, MT (500 μM), or Glutor (1 μM; glucose-transporter inhibitor) for 24 h. **(F)** Quantification of 2-NBDG mean fluorescence intensity in healthy donor- and CALRdel52het01 MPN iPSC-derived HSPC under the indicated treatment conditions. Bars show mean ± SD; n=3 biological replicates. Statistical analysis was performed via One-way ANOVA with Dunnett correction for multiple comparisons against the control group. Statistical significance is indicated by * (P < 0.05), ** (P < 0.01), *** (P < 0.001), and **** (P < 0.0001).

To functionally assess these transcriptomic data, we performed 2-NBDG glucose uptake assays. In HD progenitors, MT had no significant impact on glucose uptake compared to control, whereas in MPN iPSC-derived HSPC, MT significantly reduced 2-NBDG incorporation, comparable to the GLUT 1,2,3 inhibitor Glutor^30^ (**Fig. 2E-F**). Taken together, these data demonstrate that MT selectively impairs metabolic activity in MPN progenitors, at least in part by reducing glucose uptake, and thereby limiting bioenergetic support for malignant growth.

### Melatonin counteracts stromal activation and fibrosis-associated signaling

Given that MT selectively impairs malignant progenitors, we examined whether it also modulates stromal activation, a central driver of BM fibrosis in MPN^31^. As expected, TGF-β robustly induced alpha-smooth muscle actin (α-SMA) protein expression in primary MSC, consistent with myofibroblast transformation (**Fig. 3A-B**). MT pre-treatment markedly attenuated this response, with a significant reduction in α-SMA compared with TGF-β alone, while MT alone did not affect basal α-SMA levels. At the transcriptional level, MT suppressed TGF-β-induced induction of *ACTA2* (α-SMA)*, FN1* (fibronectin), and *GLI1*, partly normalizing their expression towards control values (**Fig. 3C**). In addition, MT impaired TGF-β-induced migratory capacity of MSC in wound-healing assays (**Fig. 3D**). These findings demonstrate that MT pre-conditioning counteracts TGF-β-driven stromal activation in monocultures.

**Figure 3.**
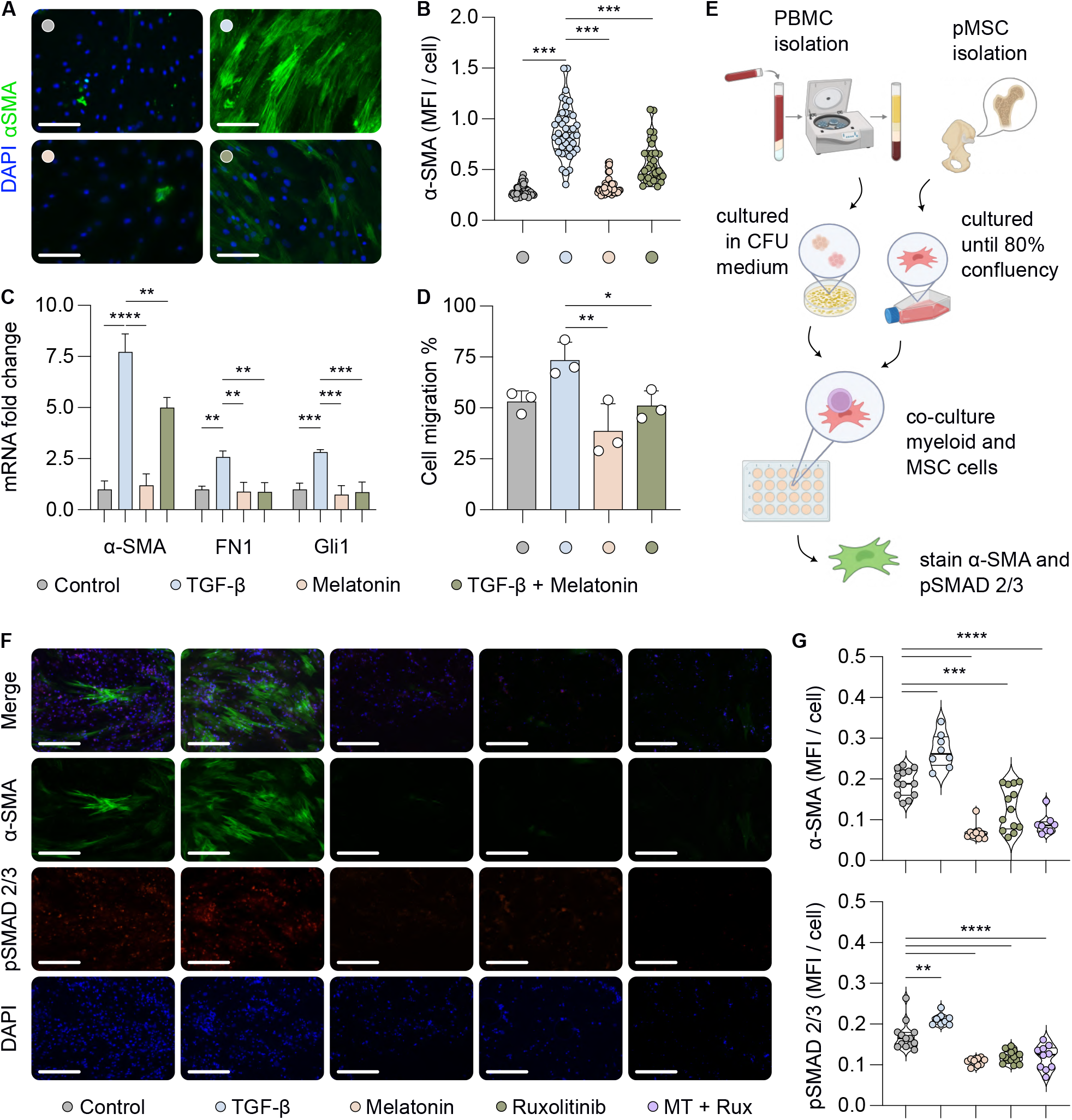
Melatonin attenuates TGF-β–driven stromal activation and MPN-associated profibrotic signaling. **(A)** Representative Immunofluorescence images of primary mesenchymal stem cells (MSC) derived from femoral heads. MSC were pre-treated with vehicle or MT (500 μM) for 48 h, and subsequently stimulated with TGF-β (10 ng/mL) for 48 h. Cells were stained for αSMA (green) and nuclei (DAPI; blue); scale bar = 125 μm. **(B)** Quantification of αSMA mean fluorescence intensity (MFI) per cell. Violin min-max plots show the median and interquartile range; n=6-8 biological replicates, n=5 technical replicates. Statistical analysis was performed via One-way ANOVA with Dunnett correction for multiple comparisons. **(C)** Reverse-transcription quantitative polymerase chain reaction (RT-qPCR) analysis of *ACTA2* (αSMA), *FN1*, and *GLI1* expression in iPSC-derived MSC co-treated with vehicle, MT (500 μM), TGF-β (2 ng/mL), or combination of MT + TGF-β for 16 h. Gene expression was normalized to *MT-ATP6* and is presented as fold-change relative to control. Bars show mean ± SD; n=3 biological replicates. Statistical analysis was performed separately for each gene via One-way ANOVA with Dunnett correction for multiple comparisons. **(D)** Cell migration in M-SOD stromal cells treated with vehicle, TGF-β (5 ng/mL), MT (500 μM), or combination of MT + TGF-β. Migration was quantified 4 h after treatment and is expressed as the percentage of wound closure. Bars show mean ± SD; n=3 biological replicates. Statistical analysis was performed via One-way ANOVA with Dunnett correction for multiple comparisons. **(E)** Schematic representation of the hematopoietic– stromal co-culture model. Myeloid cells recovered from CFU of MPN patient-derived PBMC were co-cultured with primary MSC from blood-healthy human femoral heads. After establishment of the co-culture, cells were treated for 48 h with vehicle, MT (500 μM), ruxolitinib (500 nM), or a combination of MT + Ruxolitinib. TGF-β (2 ng/mL) was included as a positive control. **(F)** Representative images of stromal cells from the co-culture experiment shown in E. Cells were stained for αSMA (green), pSMAD2/3 (red), and nuclei (DAPI; blue); scale bars = 125 μm. **(G)** Quantification of αSMA and pSMAD2/3 MFI per cell from the co-culture experiment shown in panel F. Violin min-max plots show the median and interquartile range; n=8-13 biological replicates. Statistical analysis was performed via One-way ANOVA with Dunnett correction for multiple comparisons. Statistical significance is indicated by * (P < 0.05), ** (P < 0.01), *** (P < 0.001), and **** (P < 0.0001).

To model fibrotic crosstalk between hematopoietic and stromal compartments, we employed a co-culture system of MPN patient-derived myeloid cells and primary MSC (**Fig. 3E**). Co-culture markedly induced fibrotic features, including α-SMA upregulation and activation of SMAD2/3 signaling (pSMAD2/3), which were further enhanced by TGF-β stimulation (**Fig. 3F,G**). MT treatment strongly suppressed α-SMA and pSMAD2/3 levels, comparable to the effects observed with the JAK inhibitor ruxolitinib.

### Nanoparticle delivery improves bone marrow and spleen targeting

To evaluate the therapeutic potential of MT *in vivo*, we implemented a tamoxifen-inducible SclCreER;JAK2^V617F^ mouse model^26^. Competitive BM transplants from JAK2^V617F^ or WT donors into lethally irradiated CD45.1 recipients were performed (**Fig. 4A**). High-resolution μCT imaging of long bones revealed a strong MPN phenotype, demonstrating intramedullary growth in the diaphyseal region in the recipient mice characterized by replacement of normal BM architecture with densely mineralized tissue (**Fig. 4B**). The quantitative morphological analyses revealed reduced trabecular thickness and increased diaphyseal trabecular number in JAK2^V617F^ mice compared with JAK2^WT^, reflecting abnormal bone remodeling^32^ (**Fig. 4C**).

**Figure 4.**
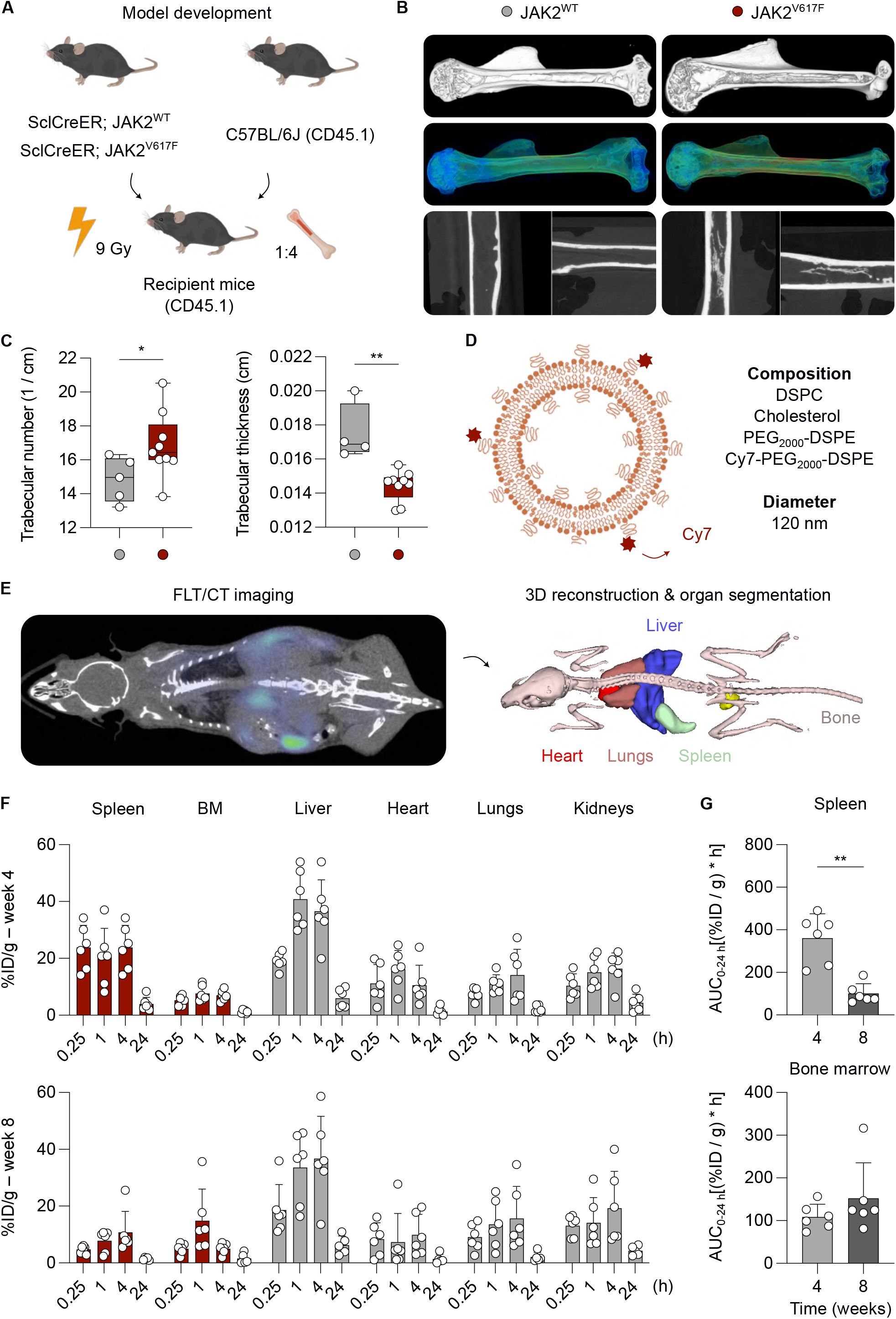
Imaging-enabled nanoparticle biodistribution assessment reveals accumulation in target organs in SclCreER;JAK2^V617F^ mice. **(A)** Schematic overview of model development. **(B)** High-resolution µCT volumetric reconstruction images indicate the development of endosteal reactions and medullary invasions along the medullary canal, contributing to a generalized osteosclerosis in the diseased JAK2^V617F^ mice; 3D bone segmentation (top), gradient map (middle), 2D coronal and sagittal views (bottom). **(C)** Bone morphometrical analysis shows the number of trabeculae at the cortical region (left) and trabecular thickness within the distal ROI (right), revealing an erratic development of the internal microarchitecture. Box-whisker min-max plots show the median and interquartile range; n=8-13 biological replicates. Statistical analysis was performed via Mann-Whitney U test. **(D)** Representation of the liposomal formulation composed of DSPC, cholesterol, and mPEG_2000_-DSPE lipids, and functionalized with Cy7-PEG_2000_-DSPE fluorescence lipids, enabling nanoparticle tracking via FLT/CT imaging. **(E)** FLT/CT image (left) and organ volumetric segmentation (right) of a mouse injected with Cy7-liposomes, visualized at 4 h post-injection. **(F)** Biodistribution profiles of the Cy7-liposomes at week 4 (top) and 8 (bottom). Each dot represents one individual mouse scanned at 0.25, 1, 4 and 24 h post-injection; n=6 biological replicates. **(G)** Comparison of the AUC of nanoparticle accumulation in the spleen (top) and the bone marrow (down) over 24 h, at week 4 versus week 8 of the disease progression. Bars show mean ± SD; n=6 biological replicates. Statistical analysis was performed via Mann-Whitney U test. Statistical significance is indicated by * (P < 0.05), ** (P < 0.01), *** (P < 0.001), and **** (P < 0.0001).

To assess the biodistribution profile of the therapeutic nano-MT in MPN mice, fluorescence tomography/computed tomography (FLT/CT) imaging of NIR dye-labeled companion diagnostic liposomes was performed at weeks 4 and 8 after transplantation (**Fig. 4D**). Upon organ segmentation, a substantial nanoparticle accumulation was observed in the target organs. In the spleen, nanoparticles accumulated in levels higher than 20% ID/g (20.3 ± 10.2 at 1 h, 23.8 ± 7.7 at 4 h, and 3.9 ± 2.3 at 24 h), while in the BM, nanoparticle targeting reached levels that are unprecedented in the case of nanoparticle-free administration (7.6 ± 2.7 at 1 h, 6.9 ± 1.8 at 4 h, and 1.2 ± 0.5 at 24 h) (**Fig. 4E,F**).

Of note, in addition to assessing the nanoparticle accumulation to the disease hot-spots at early-time points (week 4), nanoparticle accumulation was also assessed at 8 weeks post-transplantation. Interestingly, a significant drop was observed with respect to spleen accumulation at week 8 in comparison to week 4 (**Fig. 4G**). This outcome indicates temporal changes in splenic accumulation likely linked to evolving fibrosis and impaired vascularization, which may limit nanoparticle delivery or uptake despite ongoing splenomegaly.

### Liposomal melatonin remodels hematopoiesis and BM fibrosis

Mice were distributed in six comparative treatment groups aiming to assess the nano-MT therapeutic potential in MPN (**Fig. 5A**). Nanoparticle physicochemical characterization confirmed desirable characteristics, with a hydrodynamic diameter of 120 nm, narrow size distribution (polydispersity index = 0.1), and high drug encapsulation efficiency of 70% (**Fig. 5B**).

**Figure 5.**
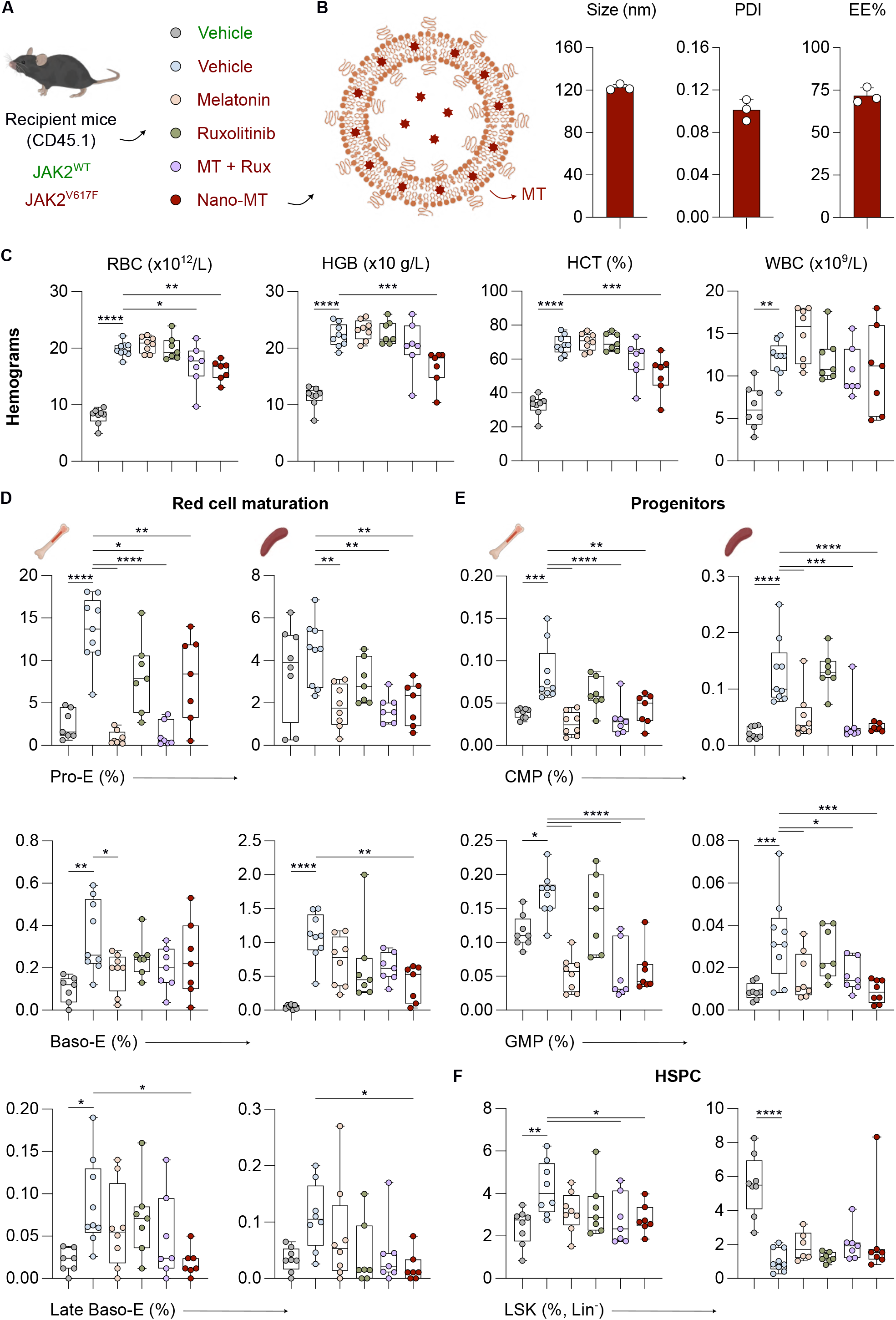
Nano-formulated melatonin suppresses erythrocytosis and abnormal hematopoietic stem and progenitor expansion in SclCreER;JAK2^V617F^ mice. **(A)** Schematic representation of the treatment strategy. Following development of a JAK2^V617F^-driven MPN phenotype, mice received vehicle, free melatonin (15 mg/kg, intraperitoneally, 5 days per week), ruxolitinib (60 mg/kg, orally, 5 days per week), a combination of free melatonin (MT) + ruxolitinib (Rux), or Nano-MT (37.5 mg/kg, intravenously, 2 days per week). JAK2^WT^ vehicle-treated mice served as controls. Treatment was administered for 6 weeks before endpoint analyses. **(B)** MT-loaded nanoparticles displayed a hydrodynamic diameter of 120 nm, low dispersity with polydispersity index (PDI) = 0.1, and 70% drug encapsulation efficiency (EE%). Bars show mean ± SD; n=3 replicates. **(C)** Peripheral blood parameters measured after 6 weeks of treatment, including red blood cell count (RBC), hemoglobin (HGB), hematocrit (HCT), and white blood cell count (WBC). **(D)** Flow cytometric analysis of erythroid maturation stages in bone marrow (left) and spleen (right) after 6 weeks of treatment, showing the frequency of pro-erythroblasts (Pro-E), basophilic erythroblasts (Baso-E), and late basophilic erythroblasts (Late Baso-E). **(E)** Frequency of myeloid progenitor populations in bone marrow (left) and spleen (right) after 6 weeks of treatment, including common myeloid progenitors (CMP) and granulocyte-monocyte progenitors (GMP). **(F)** Frequency of Lin⁻Sca-1⁺c-Kit⁺ (LSK) stem cell populations in bone marrow (left) and spleen (right) after 6 weeks of treatment. For C-F, box-whisker min-max plots show the median and interquartile range; n=7-9 biological replicates. Statistical analysis was performed via ordinary One-way ANOVA with Dunnett correction for multiple comparisons. Statistical significance is indicated by * (P < 0.05), ** (P < 0.01), *** (P < 0.001), and **** (P < 0.0001).

JAK2^V617F^ mice developed erythrocytosis and leukocytosis, with significantly elevated red blood cell (RBC) and white blood cell (WBC) counts, hemoglobin (HGB), and hematocrit (HCT) levels compared with WT mice (**Fig. 5C**). Nano-MT significantly mitigated aberrant erythropoiesis, while free MT, ruxolitinib, as well as their combination, only showed minor improvements (**Fig. 5C**).

Furthermore, erythroid maturation stages were characterized in BM and spleen via flow cytometry. In BM, JAK2^V617F^ mice displayed a significant expansion of pro-erythroblasts (Pro-E), basophilic erythroblasts (Baso-E), and late basophilic erythroblasts (Late Baso-E) in comparison to JAK2^WT^ mice (**Fig. 5D**). MT treatment significantly reduced Pro-E and Baso-E, while ruxolitinib suppressed only the Pro-E population. MT + ruxolitinib displayed comparable efficacy to free MT, while nano-MT resulted in the most effective reduction of the Pro-E and Late Baso-E subsets, approaching levels observed in JAK2^WT^ mice. Similar effects were seen in the spleen, where nano-MT attenuated extramedullary erythropoiesis, whereas other treatments showed less impact (**Fig. 5D**).

In the myeloid progenitor compartment, although ruxolitinib did not reduce the abundance of common myeloid progenitors (CMP) and granulocyte monocyte progenitors (GMP) populations, free MT, MT + ruxolitinib, and especially nano-MT pronouncedly reduced all subsets in both BM and spleen (**Fig. 5E**, **Fig. S3A-D**). Finally, nano-MT decreased the expanded Lin⁻Sca-1⁺c-Kit⁺ (LSK) population in BM, while no normalization was observed in the spleen (**Fig. 5F**).

In addition to the hematological assessment and considering the crucial role of MK in MPN disease progression, MK burden and fibrotic remodeling were evaluated in BM and spleen sections. JAK2^V617F^ mice showed marked megakaryocytic expansion. This effect was reversed upon administration of free MT, MT + ruxolitinib, and especially nano-MT. In the latter case, the MK numbers reached levels comparable to those in the JAK2^WT^ mice (**Fig. 6A,B**). The beneficial effect of nano-MT was also indicated by the strongest reduction of reticulin and collagen in comparison to all other treatment groups (**Fig. 6C,D**). Of note, upon treatment with nano-MT, reticulin and collagen expression reached levels comparable to those in healthy mice.

**Figure 6.**
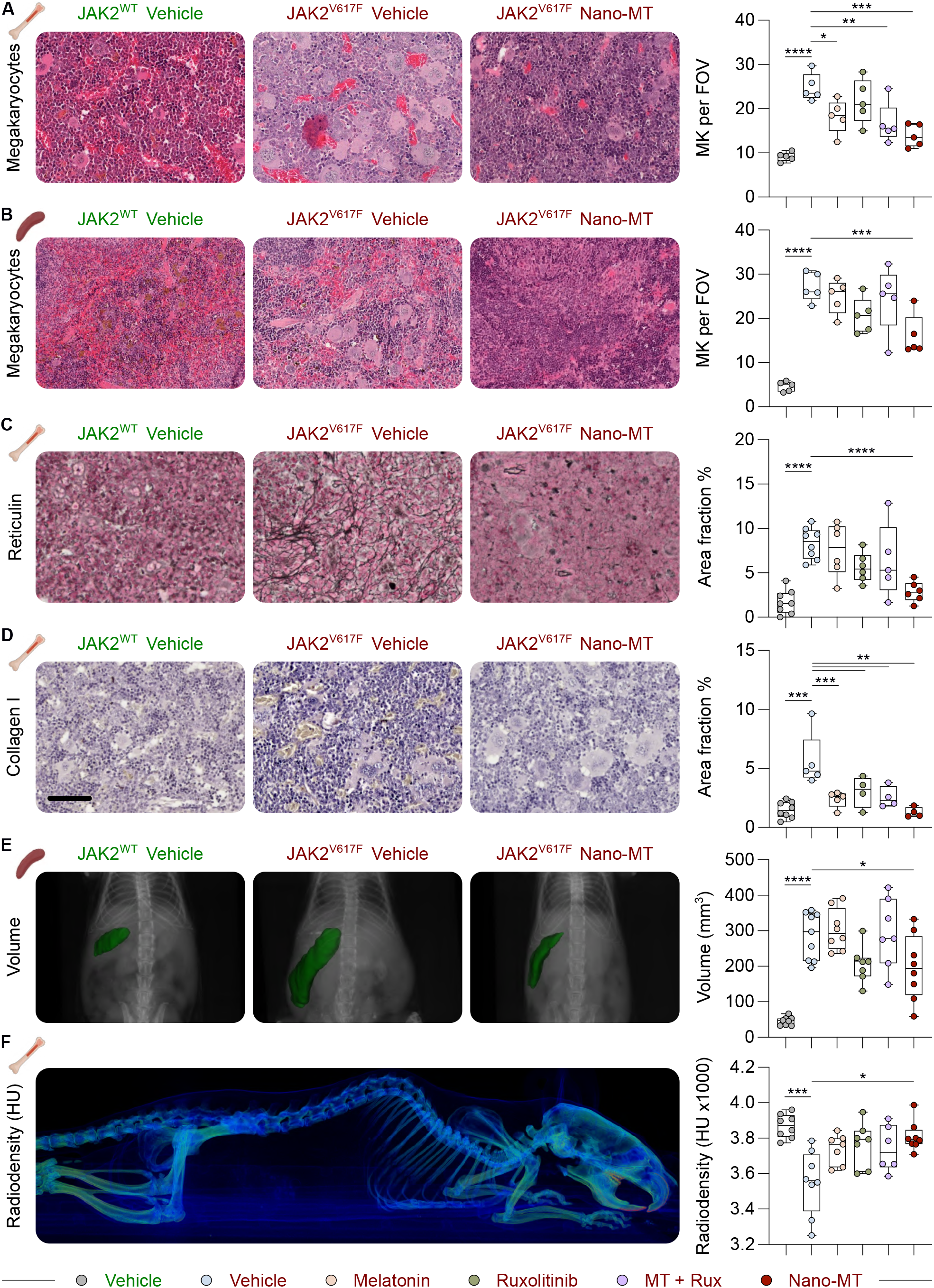
Nano-formulated melatonin reduces megakaryocyte burden and bone marrow fibrosis, and improves splenic and skeletal abnormalities in SclCreER;JAK2^V617F^ mice. **(A-D)** Hematoxylin and eosin (H&E) staining and quantification focusing on megakaryocyte (MK) number in (A) sternum and (B) spleen, as well as (C) reticulin and (D) collagen in the humeri after 6 weeks of treatment; scale bar = 200 μm. For A-D, box-whisker min-max plots show the median and interquartile range; n=7-9 biological replicates, n=5 technical replicates. Statistical analysis was performed via ordinary One-way ANOVA with Dunnett correction for multiple comparisons. **(E)** Spleen segmentation (green) and volume quantification by µCT imaging after 6 weeks of treatment / 18 weeks from model induction. **(F)** Bone radiodensity measured by µCT and expressed in Hounsfield units (HU). Segmentations were performed using fixed thresholding and further refined by manual adjustments for quality control. For E,F, box-whisker min-max plots show the median and interquartile range; n=6-9 biological replicates. Statistical analysis was performed via One-way ANOVA with Dunnett correction for multiple comparisons. Statistical significance is indicated by * (P < 0.05), ** (P < 0.01), *** (P < 0.001), and **** (P < 0.0001).

In addition, an anatomical assessment was performed. In clinical practice, splenomegaly is routinely assessed by physical examination or imaging modalities. *In vivo* µCT analysis showed significant splenomegaly in JAK2^V617F^ mice compared to JAK2^WT^ (**Fig. 6E**). The spleen volumes derived from µCT imaging significantly correlated with the terminal spleen weights at the end of the experimental procedure (**Fig. S4**). Spleen enlargement was significantly reduced following nano-MT treatment, with spleen volumes approaching those observed in JAK2^WT^ mice (**Fig. 6E**). Finally, *in vivo* radiodensity analysis of the cortical bone compartment revealed a significant reduction in Hounsfield units (HU), indicative of impaired bone remodeling and demineralization. Notably, this alteration was ameliorated exclusively by nano-MT treatment (**Fig. 6F**). Collectively, these data demonstrate that nano-MT, unlike free MT or ruxolitinib monotherapy, effectively normalizes the aberrant skeletal and hematopoietic phenotype observed in JAK2^V617F^ mice.

### Dose escalation further enhances nano-MT therapeutic efficacy

Given the large therapeutic window of MT, the nano-MT administration was intensified in an independent experiment to three times per week to maximize its therapeutic efficacy, while ruxolitinib was applied daily in single treatment and in combination with nano-MT (**Fig. 7A**).

**Figure 7.**
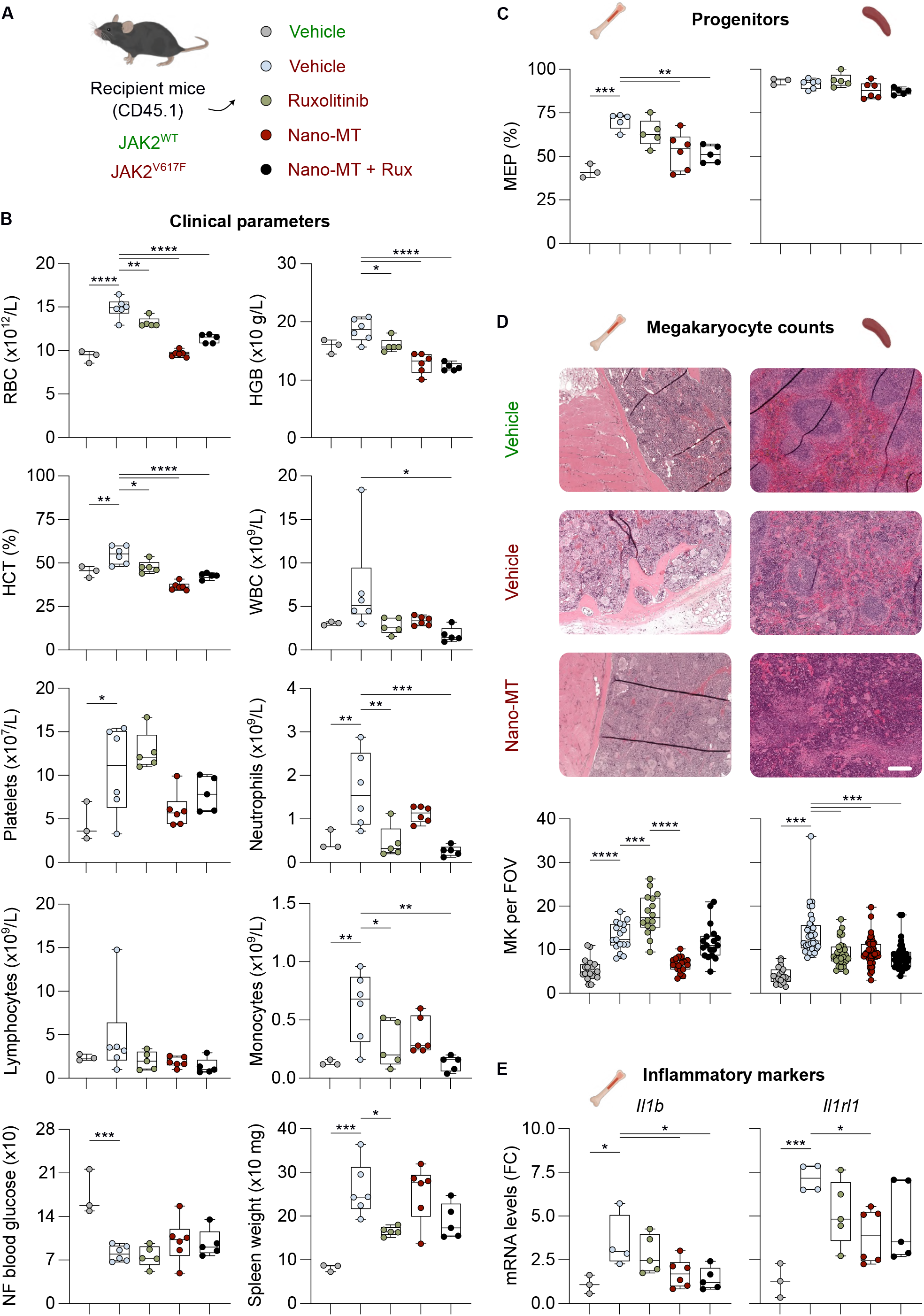
Nano-MT dose scale-up normalizes erythrocytosis, while combination with ruxolitinib improves myeloid control in SclCreER;JAK2^V617F^ mice. **(A)** Schematic representation of the treatment strategy. Following development of a JAK2^V617F^-driven MPN phenotype, mice received vehicle, ruxolitinib (60 mg/kg, orally, 7 days per week), Nano-MT (37.5 mg/kg, intravenously, 3 days per week), or a combination of Nano-MT + ruxolitinib. JAK2^WT^ vehicle-treated mice served as controls. Treatment was administered for 6 weeks before endpoint analyses. **(B)** Peripheral blood and clinical parameters measured after 6 weeks of treatment: red blood cell count (RBC), hemoglobin (HGB), hematocrit (HCT), white blood cell count (WBC), platelet, neutrophil, lymphocyte, and monocyte counts, as well as non-fasting (NF) blood glucose, and spleen weight. **(C)** Frequencies of megakaryocyte-erythroid progenitors (MEP) in bone marrow (left) and spleen (right) after 6 weeks of treatment. For B,C, box-whisker min-max plots show the median and interquartile range; n=3-6 biological replicates. **(D)** Hematoxylin and eosin (H&E) staining and quantification focusing on MK number in sternum (left) and spleen (right) after 6 weeks of treatment; scale bar = 200 μm. Megakaryocytes were quantified in at least 5 nonoverlapping fields of view (FOV) per mouse. Box-whisker min-max plots show the median and interquartile range; n=3-4 biological replicates, n=6 technical replicates. **(E)** mRNA expression levels of inflammatory markers in the bone marrow measured after 6 weeks of treatment and expressed as fold change (FC). Box-whisker min-max plots show the median and interquartile range; n=3-6 biological replicates. Statistical analysis was performed via ordinary One-way ANOVA with Dunnett correction for multiple comparisons. Statistical significance is indicated by * (P < 0.05), ** (P < 0.01), *** (P < 0.001), and **** (P < 0.0001).

JAK2^V617F^ mice displayed a robust MPN phenotype with high CD45.2 positivity (**Fig. S5**). Nano-MT monotherapy normalized erythrocytosis (RBC, HGB, HCT), reaching levels similar to those observed in JAK2^WT^ mice (**Fig. 7B**, **Fig. S6**). Ruxolitinib effects were modest compared with nano-MT alone but showed the strongest reduction in splenomegaly. The nano-MT + ruxolitinib combination yielded the strongest effects in reducing leukocytosis (WBC, neutrophils, monocytes), while platelet reduction was more pronounced in the nano-MT group, albeit not statistically significant (**Fig. 7B**). Finally, as reported previously,^33^ non-fasting blood glucose levels are significantly reduced in MPN; out of all evaluated treatment groups, only the ones containing nano-MT had a minor normalizing effect; however, due to high heterogeneity in the JAK2^WT^ group, no significant differences were detected (**Fig. 7B**). The transplantation model developed for the dose-escalation experiment revealed a pronounced bias towards the megakaryocyte-erythroid progenitor (MEP) lineage in the BM of JAK2^V617F^ mice (**Fig. 7C**). Importantly, treatment with nano-MT and nano-MT + ruxolitinib markedly reduced the proportion of MEP in the BM, whereas splenic MEP remained unaffected across treatment groups. In a similar manner, JAK2^V617F^ mice displayed a marked increase in MK counts compared to WT, which was further exacerbated by ruxolitinib monotherapy (**Fig. 7D**). Strikingly, nano-MT and nano-MT + ruxolitinib significantly reduced MK counts in the BM and spleen, with nano-MT monotherapy restoring MK levels towards physiological values. In the spleen, all treatment regimens significantly suppressed MK numbers. Finally, the fibrosis-driving markers (*IL1B* and *ST2/IL1R1L*) were ameliorated by the two groups containing nano-MT (**Fig. 7E**).

## Discussion

This study identifies MT as a dual-acting modulator of malignant hematopoiesis and stromal remodeling in MPN. Importantly, the *in vivo* application of liposomal MT formulation in SclCreER;JAK2^V617F^ mice resulted in strong therapeutic effects, e.g., normalized erythrocytosis, reduced MK burden, and ameliorated marrow fibrosis, which was not achieved with either free MT or ruxolitinib monotherapy.

Mechanistically, the selective sensitivity of MT on mutant HSPC suggests that MT modulates the dependence of the JAK2^V617F^ clone on oxidative and metabolic stress pathways. In fact, elevated ROS levels have been described as a driver of genomic instability, stem cell exhaustion, and pro-inflammatory signaling in MPN, and targeting increased ROS with N-acetylcysteine restored blood parameters in JAK2^V617F^ mice^1^. Consistent with these observations, we detected higher basal ROS levels in MPN-derived HSPC, which were significantly reduced by MT without affecting normal progenitors. This redox correction coincided with increased apoptosis and decreased colony-forming capacity, indicating that MT interferes with ROS-dependent survival signaling in the malignant clone. The inability of luzindole to reverse MT-induced apoptosis demonstrates a receptor-independent mode of action, likely mediated by direct scavenging of free radicals and stabilization of mitochondrial membranes. At the transcriptomic level, MT downregulated MYC, E2F, and G_2_/M gene sets, reflecting inhibition of proliferative, transcriptional programs that are central to JAK2^V617F^-driven expansion^19^. This pattern partly mirrors the gene expression shifts observed after IFNα therapy, where quiescence-inducing and ribosomal-suppression programs accompany eradication of mutant HSC^13,34^. However, several studies of IFNα treatment have demonstrated activation of the cell cycle and induction of differentiation, ultimately reducing long-term stem cell function, a mechanism that we have not yet investigated for MT in our study.

The connection between metabolic suppression and apoptosis in MPN progenitors is further supported by the significant reduction of 2-NBDG glucose uptake in JAK2^V617F^ cells by MT, comparable to a GLUT1/2/3 inhibitor (Glutor), suggesting inhibition of glucose transport or glycolytic flux. GSEA analyses confirmed downregulation of glycolysis, HIF1α, and DNA replication pathways, underscoring a global contraction of anabolic metabolism. These findings align with recent reports that JAK2^V617F^ HSPC rely on glycolytic reprogramming to sustain proliferation and resist oxidative stress^33,35,36^. Interestingly, we observed only a modest reduction of non-fasting blood glucose upon nano-MT treatment. As 2-NBDG fluorescence provides a cell-level surrogate readout of analogue uptake under standardized *in vitro* conditions, and non-fasting blood glucose reflects integrated whole-body glucose homeostasis, these readouts are not directly comparable^37,38^. Circulating glucose levels are influenced by feeding status, circadian regulation of hepatic glucose production, stress-dependent insulin secretion, and peripheral glucose utilization^38–40^, processes that are also modulated by melatonin signaling^41,42^. Thus, the modest, non-significant change in non-fasting blood glucose levels does not contradict the marked reduction in 2-NBDG uptake, suggesting that nano-MT preferentially affects glucose handling within the malignant hematopoietic compartment.

In addition to its effects on the mutant clone, MT exerted profound influence on the stromal compartment. *In vitro*, MT suppressed TGF-β signaling in MSC and reduced α-SMA expression and SMAD2/3 phosphorylation in MPN myeloid cell-stromal co-cultures, which restrains myofibroblast differentiation. The antifibrotic activity of MT in hepatic and pulmonary systems has been attributed to suppression of TGF-β, suggesting a conserved mechanism across tissues. However, the molecular mechanism is still unknown. In the context of MPN, where malignant MK and inflammatory cytokines perpetuate fibrotic remodeling^43,44^, the ability of MT to dampen both upstream (hematopoietic) and downstream (stromal) drivers may provide therapeutic advantage.

The translation of these fundamental findings into actual therapeutic benefit depends critically on the administration strategy. For instance, high doses of orally administered MT in mice can produce non-specific adverse effects such as motor incoordination, sedation, and behavioral and neurotoxic signs that confound interpretation of pharmacological outcomes and raise concerns about extrapolating supraphysiological rodent dosing to biologically relevant models^45,46^. Notably, we observed no behavioral side effects. Clinical trials utilizing MT in oncology have yielded inconsistent outcomes, largely due to poor oral bioavailability, rapid clearance, and variable tissue penetration^47,48^. Therefore, by capitalizing on previously optimized drug nanoformulation strategies^23^, the development and intravenous application of a liposomal MT formulation bypassed the gastrointestinal tract, increasing the overall bioavailability without raising the drug concentration, and finally direct engagement of densely packed nano-MT with malignant cell targets in PB, spleen, and BM.

Our study also highlights the value of diagnostic imaging in capturing disease progression over time^49^. To this end, high-resolution CT or MR imaging can detect BM abnormalities, osteosclerotic lesions, and the increase in fibrotic content before manifestation of systemic effects^50,51^. Furthermore, the use of imaging-based companion nanodiagnostics offers significant potential for personalized therapy by predicting nanotherapy accumulation in tissue-targets. In our study, longitudinal changes in splenic nanoparticle accumulation between the early (week 4) and late (week 8) stages of disease reflect a functional transition to a fibrotic phenotype. Clinical implementation of such companion nanodiagnostic strategies could provide a surrogate marker of therapeutic response and improve patient stratification by identifying individuals with high nanoparticle accumulation for treatment or clinical trial inclusion, while excluding those with low accumulation, thereby maximizing therapeutic benefit.

From a therapeutic point of view, the high nano-MT accumulation in hematopoietic organs explains the superior results compared to free MT. These data underscore that previous failures of MT in oncological applications can be associated with pharmacological limitations rather than a lack of biological potential. By enhancing delivery accuracy and stability, the liposomal formulation appears to unlock MT’s dual cytoreductive and microenvironmental homeostatic effects. Nano-MT revealed distinct mechanisms when compared to established therapies. IFNα can induce molecular remission in subsets of patients by targeting quiescent HSC^13^, and recent data demonstrate that also ruxolitinib reduces mutant variant allele frequency in MPN^12^. In our model, nano-MT combined with ruxolitinib at suboptimal dosage yielded additive hematologic improvements and further suppression of inflammatory mediators such as IL-1β^9^ and IL1RL1/ST2^52^. Hence, MT may enable dose reduction of ruxolitinib in the clinical setting, potentially minimizing treatment-related adverse effects.

While the translational trajectory of such a strategy is promising, several aspects warrant careful evaluation^53^. The long-term impact of sustained antioxidant exposure on normal hematopoiesis remains uncertain, as physiological ROS fluctuations are integral to stem cell signaling^54^. It will be essential to define a therapeutic window that minimizes malignant proliferation without compromising normal HSC function. Pharmacokinetic and safety studies in animal models, followed by clinical evaluation, as exemplified by N-acetylcysteine (NCT05123365)^55^, will be required to establish optimal dosing regimens and assess potential off-target effects. Nonetheless, the favorable safety record of MT in other clinical contexts provides reassurance for early-phase testing. Consistent with our findings, MT nanoformulations have previously been assessed for their *in vivo* safety, improvement of bioavailability, and robust therapeutic efficacy, supporting the translational potential of such nanoparticle-based delivery strategies^25,56^.

Reflecting on the increasing focus on disease-modification, several promising therapeutic agents are currently under investigation for MF. These include molecules in late-stage development, such as: pelabresib^57^, a BET inhibitor; navtemadlin (KRT-232)^58^, an MDM2 inhibitor; selinexor^59^, an XPO1 inhibitor; imetelstat^60^, a telomerase inhibitor; INCA033989^61^ and JNJ88549968^62^, mutant CALR-targeting antibodies currently in early-phase clinical trials. Given MT’s broad pleiotropic properties, including antioxidant, anti-inflammatory, and immunomodulatory activities, MT and analogues may represent complementary homeostatic agents that could be explored in combination with targeted therapies to enhance overall disease management.

Taken together, this work identifies nano-MT as a pharmacologically pleiotropic agent with potency against JAK2^V617F^ MPN, targeting the malignant clone and its niche, and exploits metabolic and microenvironmental dependencies that sustain MPN pathogenesis. Given the limitations of existing treatments and the accessibility of MT-based formulations, clinical exploration of nano-MT represents a rational and potentially transformative step in MPN management.

## Acknowledgements

The authors gratefully acknowledge financial support from the German Research Foundation (DFG)- funded Clinical Research Unit 344 to NC (CH1509/1-1; 417911533), AMS (SO2198/1-2), and SKos (KO2155/7-1; 428858786); the Excellence Initiative RWTH JPI 2021 to AMS; Research Training Group 2375 (331065168) to SE, JL, FK, TL, and FDL; research grant 537070058 to JB; SFB TRR219 (322900939; subproject M07) and the Corona Foundation (S199/10084/2021) to E.P.C.v.d.V.; and the German Cancer Aid (SDK; Postgraduate Program MSSOABCD) to AMS, SKos, JB, FDL, and TL. This work was supported by the Flow Cytometry and Immunohistochemistry Facility, Core Facilities of the Interdisciplinary Center for Clinical Research (IZKF) within the Faculty of Medicine at RWTH Aachen University (RRID:SCR_028757). Biomaterial samples were provided by the RWTH centralized Biomaterial Bank Aachen (RWTH cBMB, Aachen, Germany) in accordance with the regulations of the biomaterial bank and the approval of the Ethics Committee of the Faculty of Medicine, RWTH Aachen University. The JAK2V617F knock-in mouse model was provided by the Institute Gustave-Roussy and Isabelle Plo.

## Author contribution

NC and AMS conceptualized and supervised the study. SG, AM, MAST, AMS, and NC designed the experiments. SKos collected patient samples and provided clinical data. SE, SKho, and AMS developed the nanoformulations. KO and MAST developed and provided the iPSC lines. SG, VS, KP and NC performed the *in vitro* studies. SG, AM, AN, FK, TL, FDL, JB, AMS, and NC performed the *in vivo* studies. MV, JL, MJR, PW, VH, CBL, CZ, HJ, BJ processed mouse organs for FACS and other analyses. AM performed imaging analysis. TNR and E.P.C.v.d.V. provided support for data analyses. SG, AM, AN, AB, and FDL performed histological analysis. SG and AM analyzed the data and performed statistical analysis. SG, AM, AMS, and NC designed the figures. SG, AM, AMS, and NC drafted the first version of the manuscript. All authors read, corrected, and approved the final version of the manuscript.

## Additional information

The online version contains supplementary material.

