## Supplementary Data for "Melatonin nanoparticles inhibit mutant hematopoiesis & restore bone marrow architecture in myeloproliferative neoplasms"

### Supplementary Methods

#### Isolation of peripheral blood mononuclear cells (PBMC) and mesenchymal stromal cells (MSC)

PBMC were isolated from peripheral blood of MPN patients/healthy donors (**Table S1**) by density-gradient centrifugation (Pancoll, Pan Biotech, Aidenbach, Germany). After red cell lysis, (Erylysis-Buffer pH 7.2 - 7.4 (sterilized), Morphisto Ltd, Offenbach am Main, Germany), the cells were counted using CASY cell counter (Omni Life Sciences, Bremen, Germany) and subjected to different assays described in this manuscript. All MPN samples were collected at the Department of Hematology, Oncology, Hemostaseology and Stem Cell Transplantation at RWTH Aachen University Medical Center after patients' written informed consent, as approved by the local ethics committee (EK 127/12). Primary MSC were isolated from femoral heads as described before[1] and HD samples were provided by the Transfusion Medicine in the University Hospital, RWTH Aachen (EK 099/14, EK 300/13).

#### Colony Formation Unit (CFU) assay

After isolating PBMC from primary MPN/healthy donor (HD) samples, cells were plated at a density of $10^6$ cells/mL in methylcellulose (MethoCult™, StemCell Technologies Vancouver, Canada) with 20 mL IMDM (Gibco, Massachusetts, USA), 50 ng/mL hSCF, 10 ng/mL hIL-3, 10 ng/mL hGM-CSF, and 14 ng/mL hEPO (all from Immunotools, Friesoythe, Germany) supplemented with either vehicle (0.25% DMSO) or melatonin (MT) at concentrations of 250, 500, and 750 µM. After 14 days, total colony numbers and cell counts were evaluated.

For $CD34^+$ enriched induced pluripotent stem cell (iPSC)-derived MPN/HD HSPC, 5000 cells/mL were seeded in triplicate into 24-well plates. Treatments included vehicle (0.25% DMSO), MT (500 µM, Sigma Aldrich, Missouri, USA), IFNα (1000 U/mL, Immunotools), or a combination of these agents.

#### Cultivation of M-SOD cells

The human h-tert-$EGFP^+$ mesenchymal stromal cell line (M-SOD) was kindly gifted by the laboratory of Ivan Martin, Department of Biomedicine, Basel University Hospital, Basel, Switzerland. The cells were cultured as described previously[2].

#### Apoptosis assay

Apoptosis was measured using the Pacific Blue™ Annexin V Apoptosis Detection Kit with 7-AAD (BioLegend, San Diego, USA) following the manufacturer's protocol. iPSC-derived MPN/HD hematopoietic stem and progenitor cells (HSPCs) were treated for 72 h with vehicle (0.25% DMSO), MT (500 µM), luzindole (10 µM) or combination. Afterwards, the cells were collected, washed twice with PBS/2% FCS, centrifuged at 300g without brakes (acc: 5, dec: 2) and resuspended in Annexin

V Binding Buffer (0.25-1X$10^6$ cells/mL). 100 µL of cell suspension was combined with Pacific Blue annexin V (1 µL/test) and 7-AAD (10 µL/test) in flow cytometry tubes, incubated for 10–15 minutes at room temperature in the dark, then diluted with 400 µL Binding Buffer. Samples were analyzed by flow cytometry (BD FACS Canto II) within 1 hour. Data were analysed using FlowJo v10 (BD Biosciences, New Jersey, USA) and viable (Annexin $V^-$/7-$AAD^-$), early apoptotic (Annexin $V^+$/7-$AAD^-$), late apoptotic/secondary necrotic (Annexin $V^+$/7-$AAD^+$), and necrotic (Annexin $V^-$/7-$AAD^+$) populations were identified.

**PKH-dye based mixed culture competitive apoptosis assay**

PV patient- or HD-derived $CD34^+$ HSPC were seeded at 10,000 cells per well (24-well format) and pre-cultured for 48 h in StemSpan™ SFEM (StemCell Technologies) supplemented with 100 ng/mL SCF, 100 ng/mL FLT-3L, 20 ng/mL IL-3 (all Immunotools, Friesoythe, Germany) and 20 ng/mL hyper–IL-6 (kind gift from Stefan Rose-John, Department of Biochemistry, Kiel, Germany). Subsequently, the cells were harvested and labelled with PKH dye (PKH67, HD or PKH26, PV) as per the manufacturer's protocol (Thermo Fisher, Massachusetts, USA). The labelled cells were mixed in 1:1 ratio, treated with vehicle (0.25% DMSO) or MT (500 µM) and incubated for 72 h. HD and PV populations were discriminated by $PKH67^+$ and $PKH26^+$ gates respectively, and the frequency of apoptotic cells was quantified as Annexin $V^+$ within each PKH labelled population upon treatment by flow cytometry (BD FACS Canto II).

**Reactive oxygen species (ROS) detection assay**

iPSC derived MPN/HD $CD34^+$ MACS-enriched HSPC were treated with vehicle (0.25% DMSO), MT (500 µM), Luz (10 µM) or combination in HSPC progenitor medium for 72 h. The medium composition is presented in **Table S6**. After 72 h, the cells were centrifuged at 300 g, 4 °C, and supernatant was discarded. Around 500,000 cells/condition were stained with 25 µM dihydroethidium (DHE, AAT Bioquest®, Pleasanton, USA) for 30 min at 37 °C in the dark and ROS levels were detected in PE channel by flow cytometry.

**3'mRNA-Seq**

6x MPN- (3x PV, 3 MF) and 3x HD-derived $CD34^+$ HSPC (**Table S1**) were treated with vehicle (0.25% DMSO), or MT (500 µM) for 16 h in HSPC progenitor medium supplemented with 10 ng/mL SCF, 10 ng/mL FLT-3L, 2 ng/mL IL-3 and 2 ng/mL hyper-IL6. Total RNA was isolated using RNeasy Micro Kit (Qiagen, Venlo, Netherlands) according to the manufacturer's instructions, including on-column DNase digestion. RNA quantity and integrity were assessed and samples meeting quality thresholds were processed for library preparation. 3′ mRNA-seq libraries were prepared using the QuantSeq 3′ mRNA-Seq Library Preparation Kit FWD for Illumina (Lexogen, San Diego, USA) following the manufacturer's protocol. Sequencing was performed by Genomics Facility of the Interdisciplinary Center for Clinical Research (IZKF) Aachen within the Faculty of Medicine at RWTH Aachen

University on an Illumina NextSeq 500 platform using single-end reads. Each sample yielded an average of ~7.66 million reads per library. The data generated in this study have been deposited in the Gene Expression Omnibus (GEO) under accession number GSEXXXXXX.

**Bioinformatics**

Raw sequencing data were processed using a standard RNA-seq workflow. Read quality was assessed with FastQC. Reads were aligned to the human reference genome [GRCh38/hg38]. Differential gene expression analysis (DGEA) was performed in Rstudio (Posit, Massachusetts, USA) using DESeq2 applying the Benjamini-Hochberg procedure to control the false discovery rate (FDR). Genes with adjusted p-value < 0.05 and $\log_2 FC$ > 1.5 fold were considered significant. Volcano plots and pathway enrichment were performed using ggplot2 based on the differentially expressed genes (DEG) list.

Gene Set Enrichment Analysis (GSEA) was performed using the Broad Institute GSEA software (version 4.3.3). Human Molecular Signatures Database (MSigDB) collection (H: Hallmark) was utilized for the GSEA analysis. Enrichment scores were calculated using the weighted Kolmogorov-Smirnov-like statistic, and significance was assessed using permutation testing (gene set permutations; n = 1000). Normalized enrichment scores (NES) and FDR q-values are reported; gene sets with FDR q < 0.05 were considered significantly enriched.

**Glucose uptake assay**

iPSC-derived MPN/HD HSPC were cultured in progenitor medium (**Table S6**) supplemented with 100 ng/mL SCF, 100 ng/mL FLT-3L, 20 ng/mL IL-3, and 20 ng/mL hyper–IL-6 for 48 h, followed by treatment with vehicle (0.25% DMSO), MT (500 μM), or Glutor (1 μM, MedChemExpress, New Jersey, USA). Following treatment, cells were incubated for 2 h in glucose-free medium (DMEM, no glucose, Gibco) supplemented with 0.5% human serum albumin (HSA, Grifolls). Cells were subsequently incubated with 2-NBDG (50 μM, Thermo Fisher Scientific) for 1 h at 37°C protected from light. Uptake was terminated by flow cytometry. Glucose uptake was quantified as the mean fluorescence intensity (MFI) of 2-NBDG in the analyzed population, and representative histograms were displayed normalized to mode.

**Monoculture – Immunofluorescence staining**

Primary MSC (25,000 cells) were seeded on coverslips previously coated with 0.1% gelatin (Sigma Aldrich) in DMEM, low glucose (Thermo Fisher) with 2% FCS. After 24 h, the cells were pre-treated with vehicle (0.25% DMSO) or MT (500 μM) for 48 h. Afterwards, the medium was refreshed and subsequently, TGF-β (2 ng/mL) was added to induce α-SMA expression. The fibrotic cells were fixed and stained as per manufacturer's protocol[3]. A list of antibodies used can be found in **Table S4**.

α-SMA signal was quantified in ImageJ (NIH, USA) by segmenting nuclei (DAPI) to estimate cell number and measuring α-SMA MFI per image. Background was subtracted using the rolling-ball radius option. α-SMA intensity was normalized to cell count to obtain MFI per cell. For each condition, at least 5 fields were analyzed across 3 biological replicates and data were plotted as violin distributions.

**iPSC-derived mesenchymal stromal cell differentiation, culture and RT-qPCR**

iPSC-derived mesenchymal stromal cells (iMSCs) were generated using a modified embryoid body (EB)-based differentiation protocol as previously described[4]. Briefly, iPSC-derived EBs were transferred to 0.1% gelatin-coated six-well plates (Sigma) to promote EB attachment and cultured in low-glucose DMEM (Gibco) supplemented with 10% fetal calf serum (FCS; PAN-Biotech), 10 ng/mL basic fibroblast growth factor (bFGF; ImmunoTools), and 10 ng/mL bone morphogenetic protein 4 (BMP4; Miltenyi Biotec) for 4 days. From day 5 onward, the cells were maintained in the same culture medium without BMP4. For the first three passages, cells were detached using Accutase (PAN-Biotech) and transferred to 0.1% gelatin-coated 10-cm culture dishes. From passage 4 onward, cells were cultured on uncoated tissue-culture plastic. iMSCs between passages 4 and 8 were used for all experiments. For quantitative RT-qPCR, WT iMSC (80,000 cells/well) were seeded on a six well plate and allowed to attach for 24 h. Afterwards the cells were pre-treated with 0.25% DMSO or MT (500 µM) for 30 mins. The medium was refreshed followed by 48 h stimulation with TGF-β (10 ng/mL). Total RNA was isolated as per the manufacturer's protocol (GeneJET RNA Purification Kit, Thermo Fisher). cDNA was synthesized using 500 ng total RNA and gene expression was analyzed using SYBR™ Select Mastermix (Thermo Fisher) on TaqMan 7500 Fast device (Applied Biosystems). A list of primers used can be found in **Table S2**.

**Co-culture assay**

PBMC derived from MPN patients were subjected to CFU assay for 14 days as described previously. On day 12, MSC were seeded on gelatin-coated coverslips for 48 h until 70-80% confluent. Afterwards, the CFU colonies were harvested gently by diluting the methyl cellulose with PBS-2%FCS. The harvested cells were centrifuged at 300 g for 10 mins, washed and resuspended in 1 mL RPMI1640 (Pan Biotech) supplemented with 10% FCS. The cells were mixed with 0.4% trypan blue solution (1:1) and counted using the improved Neubauer hemocytometer (4 large squares). Total 500,000 cells per condition in 300 µL RPMI1640/10% FCS were seeded on top of MSC. The co-culture was performed for 7 days. On day 5, the co-culture was treated with the following compounds, co-culture (0.25% DMSO), TGF-β (2 ng/mL, Immunotools), MT (500 µM, Sigma Aldrich, Missouri, USA), ruxolitinib (500 nM, Selleck Chemicals GmbH, Munich, Germany), and combination of MT + ruxolitinib for 48 h. The cells were fixed and stained as described above. A list of antibodies used can be found in **Table S5**.

### Scratch Assays

M-SOD cells (10,000 cells/0.5 mL) were seeded in 24-well plates (Greiner) for 24 h. When the cells achieved 70% confluency, a linear scratch was made using a pipette tip (200 µL) followed by the cells were treated with vehicle (0.25% DMSO), TGF-β (5 ng/mL), MT (500 µM) and combination for 4 h. Afterwards, the cells were imaged in GFP channel at EVOS M7000 (Thermo Fisher) at 10x magnification. The wound closure/cell migration was quantified using ImageJ as described[5].

### Animal experiments

C57BL/6J SclCreER;JAK2$^{V617F}$ KI[6,7] (CD45.2$^{+}$), C57BL/6J SclCreER;JAK2$^{WT}$ (CD45.2$^{+}$; littermate controls), and C57BL/6J CD45.1 (rescue bone marrow donors and transplant recipients) were used. Both male and female animals were included. Donor mice were 8-14 weeks old; recipient mice were 10–20 weeks old. All lines were bred in-house at the RWTH Aachen animal facility. Following transplantation, recipients were housed for the first 14 days at the Institut für Versuchstierkunde (VTK) and then moved to the closed barrier system at the Institute for Experimental Molecular Imaging (ExMI) for continued housing and *in vivo* imaging. To induce Cre-mediated activation of JAK2$^{V617F}$, tamoxifen (100 µg/g body weight) was administered intraperitoneally (i.p.) once daily for 5 consecutive days. Donor mice were monitored by peripheral blood sampling (tail vein; ~40 µL) beginning 6 weeks after tamoxifen. Recipient WT C57BL/6J CD45.1 mice received a single 9 Gy lethal irradiation dose. Within 24 h of irradiation, recipients were transplanted via lateral tail vein injection with a mixture of $2\times10^{6}$ (1$^{st}$ experiment) or $1.5\times10^{6}$ (2$^{nd}$ experiment) BM cells in PBS from SclCreER;JAK2$^{V617F}$ (or SclCreER;JAK2$^{WT}$) donors plus $1\times10^{6}$ (1$^{st}$ experiment)or $0.5\times10^{6}$ (2$^{nd}$ experiment) rescue BM cells from WT CD45.1 donors. To reduce infection risk during engraftment, cotrimoxazole (100 µg/mL) was provided in drinking water for 2 weeks post-transplant. Engraftment and disease phenotype were monitored by serial tail-vein blood sampling, with analysis by flow cytometry (e.g., CD45.1 vs CD45.2, and lineage markers such as Gr1/CD11b and Ter119).

Treatment was initiated 12 weeks post-transplantation, when BM fibrosis grade 1–2 was expected. Groups were: (i) JAK2$^{WT}$ vehicle; (ii) JAK2$^{V617F}$ vehicle; (iii) JAK2$^{V617F}$ free-MT; (iv) JAK2V617F Ruxolitinib; (v) JAK2V617F free-MT + ruxolitinib; (vi) JAK2V617F nano-melatonin (nano-MT); and (vii) Cy7-liposome (nano-Cy7) for biodistribution imaging. In the dose escalation experiment, the groups were streamlined to (i) JAK2$^{WT}$ vehicle; (ii) JAK2$^{V617F}$ vehicle; (iii) JAK2$^{V617F}$ ruxolitinib (iv) JAK2V617F nano-MT; (v) JAK2V617F nano-MT + ruxolitinib.

For procedures requiring anesthesia (injections requiring immobilization and all imaging), mice received inhalational isoflurane (induction 5 vol% in oxygen in an induction chamber; maintenance 2% vol via face mask). Eyes were protected with Bepanthen (Bayer AG, Leverkusen, Germany) eye/nasal ointment.

**Drug formulation and dosing**

Free MT (#Cat. 444300, Sigma Aldrich) was prepared by dissolving the compound in PBS with 1% DMSO. The compound (15 mg/kg) was administered by i.p. injection, 5 days/week.

Ruxolitinib phosphate (#Cat. HY-50858, MedChem Express) was formulated by dissolving the compound in 0.5% (w/v) hydroxypropyl methylcellulose (Sigma Aldrich). Mice received 60 mg/kg ruxolitinib by oral gavage (5 days/week or 7 days/week), with the dosing volume adjusted to body weight. For the combination therapy (MT + ruxolitinib), the compounds were administered sequentially with the same dosage and routes as described previously for 5 days/week.

**Nano-formulation synthesis and dosing**

Nano-MT was prepared using a microfluidic method[8]. Briefly, a lipid solution was prepared by dissolving DSPC 18:0 PC (1,2-distearoyl-sn-glycero-3-phosphocholine; Avanti® Polar Lipids, Merck, Darmstadt, Germany), cholesterol (Sigma Aldrich), and methoxy-PEG2000-DSPE (Lipoid GmbH, Ludwigshafen, Germany) in ethanol at a molar ratio of 62:33:5, corresponding to a final liposome concentration of 30 mM. MT was incorporated into the lipid phase, to a corresponding final concentration of 6 mg/mL in the formulation. Both lipid solution and aqueous solution (PBS) were preheated to 45 °C and transferred into polypropylene syringes mounted on an advanced programmable syringe pump (Harvard Apparatus PHD ULTRA, Holliston, USA). Nano-MT was formed by mixing the two phases in a T-junction mixer at a flow rate ratio of 2:1 (aqueous: organic) and a total flow rate of 20 mL/min. The resulting Nano-MT liposome was maintained on a hot plate at 70 °C under stirring until the organic solvent was completely evaporated.

The average particle size and polydispersity index (PDI) were determined by dynamic light scattering (DLS; Malvern Instruments Ltd., UK). MT concentration was quantified by absorption spectroscopy following a previously described method[9]. In brief, samples were disrupted by dilution in isopropanol (1:40, v/v) and vortexed for at least 30 s to ensure complete liposome disassociation and MT release. Subsequently, 100 µL of disrupted Nano-MT was compared to the MT standard curve (8 concentrations; 0–0.4 mg/mL of MT concentration) prepared in isopropanol containing the same lipid concentration as the samples, to correct for background absorbance. Absorbance was measured at 300 nm using a TECAN Infinite M200 PRO microplate reader (Tecan Trading AG, Switzerland). Encapsulation efficiency (EE%) was calculated according to the following equation: EE% = (Measured MT concentration / initial MT concentration added during preparation) *100. For each injection time point, the formulation was freshly prepared and subsequently concentrated to match the desired injected dose per kg of mouse body weight. The compound was administered 2 days/week or 3 days/week by i.v. injection into the tail vein.

**In vivo biodistribution analysis**

The *in vivo* biodistribution of the Cy7-labeled Nano-MT was investigated using a hybrid micro-computed tomography-fluorescence tomography (FLT/CT) device (U-CT OI, MILabs B.V., Utrecht, the Netherlands) at 0.25, 1, 4, and 24 h, to emphasize on the early/late time points post-injection. LP were synthesized as described before, with 0.2 mol% of Cy7-PE added to the organic phase before mixing. For each administration, a lipid concentration of 7 mM was used. Utilizing a dedicated vaporizer (Harvard apparatus, Holliston, USA), anesthesia was induced during all the in vivo experiments using 5% isoflurane (Forene, Abbott, Wiesbaden, Germany). During the imaging process, isoflurane concentration was reduced and maintained at 2%, and eyes were properly covered with Bepanthen eye ointment (Bayer Vital GmbH, Leverkusen, Germany) to prevent desiccation. Mouse temperature was kept constant via use of a heating pad. Labelled NP were injected into the tail vein of the mice using a sterile catheter consisting of a 30 G cannula (B. Braun, Melsungen, Germany) connected to a polyethylene tube (Hartenstein, Würzburg, Germany). For imaging, anesthetized mice were placed in a dedicated animal holder and inserted in the optical unit of the FLT/CT (laser/filter system with excitation and emission wavelengths at 730 and 775 nm, respectively, acquiring 130 scan points). Subsequently, the animal holder moved to the CT automatically, acquiring a total body scan (480 projections 1944 × 1536 pixels, full rotation in step-and-shoot mode, 55 kV, 0.17 mA, 75 ms). The process was repeated for each mouse at 0.25, 1, 4, and 24 h post-injection after 5 and 7 weeks from disease induction to visualize the fluorescence signal. After the last FLT/CT scan, the animals were sacrificed by cervical dislocation under anesthesia. The FLT/CT images, shape, scattering maps, and absorption maps were automatically reconstructed[8]. The FLT/CT data were used to construct 3D organ segmentations using interactive segmentation procedures (Imalytics Preclinical, Gremse-IT GmbH, Aachen, Germany) with a previously developed technique[8]. Organs of interest were volumetrically segmented for all mice and all time points. The total body fluorescence signal from the FLT scans at 0.25 h post-injection was used for the calculation of total NP concentration (% injected dose per gram – %ID/g)[8]. Means and standard deviation were calculated based on six mice for each time point for both weeks 10 and 7. The area under the curve (AUC) for both weeks 4 and 8 was calculated to compare the distribution of NP in the organs.

**Micro-computed tomography (µCT)**

*In vivo* micro–computed tomography (µCT) scans were performed using a high-resolution preclinical scanner (MILABS, Utrecht, The Netherlands). Animals were anesthetized and positioned in the scanner to minimize motion during image acquisition. µCT scans were acquired using a tube voltage ranging from 20 to 65 kVp and a tube current of 0.17 mA, with an exposure time of 75 ms per projection. A combined aluminum filter (100 + 400 µm) was applied to optimize beam quality and reduce beam-hardening artifacts. Images were acquired with an isotropic voxel size of 0.04 mm, resulting in a field of view of 64 × 64 × 100 mm. Data were collected over a full rotation using a 0.25° rotation step in accurate scanning mode. The estimated radiation dose for the in vivo protocol was approximately 410 mGy. Image reconstruction was carried out using the manufacturer’s software with standard correction algorithms for beam hardening and ring artifacts. Reconstructed datasets were used for qualitative and quantitative analyses. Quantitative morphometric analyses were conducted using a dedicated software (Imalytics Preclinical, Gremse-IT GmbH, Aachen, Germany). Briefly, bone regions of interest (ROIs) were delineated by using a fixed-threshold approach. A threshold of 2000 Hounsfield Units (HU) was applied to selectively segment mineralized bone tissue, ensuring consistent and reproducible ROI definition across all specimens. To ensure precise measurements, calibration phantoms containing distilled water and two 4 mm Bruker rods with densities of 0.3 and 1.25 g/cm³ CaHA were used and routinely checked for consistency. Cortical bone ranged between 3000-4000 HU, corresponding to 485-616 mgHA/cm³.

**Histopathology**

Freshly collected organs were processed and stored as follows. Spleens were weighed and sectioned into three portions. The first portion was embedded in Tissue-Tek® O.C.T.™ Compound (Sakura Finetek Europe B.V., Alphen aan den Rijn, The Netherlands), snap-frozen in liquid nitrogen, and stored at −80 °C. Cryosections (8 µm) were prepared using a Cryostat CM3050 S (Leica, Wetzlar, Germany). The second spleen portion was fixed in 4% paraformaldehyde (PFA), while the third portion was used for tissue disaggregation and flow cytometry (FACS) analysis. Long bones were carefully cleaned of surrounding tissue prior to processing. Femurs were used for bone marrow isolation followed by disaggregation and downstream analyses. Humeri were fixed in 4% PFA, placed in plastic tubes, stabilized with styrofoam to prevent movement, and tightly sealed to avoid dehydration and motion artifacts during rotation. After 48 h of fixation, PFA was replaced with 70% ethanol, and samples were stored at 4 °C until scanning. The contralateral bones, and sterna were processed for decalcification following manufacturer’s protocol; briefly, bones were fixed in 4% PFA for at least 72 hours, gently washed with PBS three times, and transferred to a 25% 7.4 pH EDTA solution (Morphisto GmbH, Frankfurt am Main, Germany). Bones were transferred to plastic tubes at 4° on a moving plate for up to 10 days, the decalcifying solution was changed every 48 hs. Finally, samples were transferred in 70% EtOH, and submitted to the Immunohistochemistry Facility of the Interdisciplinary Center for Clinical Research (IZKF) within the Faculty of Medicine at RWTH Aachen University for routine processing, paraffin embedding, sectioning, and hematoxylin and eosin (H&E) staining.

For H&E staining, samples were dehydrated, embedded in paraffin, and sectioned at 5 µm using a SLIDE4003E microtome (pfm Medical, Cologne, Germany). The sections were fixed on adhesive microscope slides, deparaffinized, and stained using an automated slide-staining station (Thermo Fisher). The slides were stained in hematoxylin for 5–10 min; rinsed in warm water for 10 min; stained in 0.3% eosin for 5 min; rinsed again with tap water; and successively dehydrated in 70%, 96%, and 100% ethanol. Then, the slides were treated with xylene and sealed with glass coverslips using Vitro-Clud® (Langenbrink GmbH, Emmendingen, Germany). Image acquisition was performed using a digital slide scanner (Fritz Precipoint, Garching bei München, Germany).

**Megakaryocyte quantification**

Megakaryocyte (MK) numbers were quantified on H&E–stained sections from spleens and sterna (bone marrow). Slides were digitized and analyzed using ViewPoint Light software (Precipoint, Garching, Germany). MKs were identified based on characteristic morphology (large cell size with abundant eosinophilic cytoplasm and multilobulated/hyperchromatic nuclei) and counted within randomly selected, non-overlapping high-power fields per section. For each mouse, ≥5 fields were quantified per tissue, avoiding areas with cutting artifacts or tissue folds. MK counts were expressed as MKs per field for each animal, with the mean per mouse used for group-level comparisons.

**Reticulin staining and quantification**

Reticulin fiber staining in spleen and bone marrow sections was performed using a commercially available kit (Sigma-Aldrich) according to the manufacturer's instructions. Quantification of reticulin fibers was carried out using an image analysis approach adapted from a previously published method[10]. Briefly, 20x magnification images were exported and analyzed using ImageJ. Manual color thresholding was applied to identify the reticulin network, followed by isolation of the highlighted mesh using shape-based filtering. The Shape Filter plugin was applied with the following parameters: area 0.001–∞, Feret diameter 0.05–∞, circularity 15–∞, elongation 0.05–1, and solidity 0.5–1, with black background and "draw holes" options enabled. This workflow allowed consistent visualization and quantification of the reticulin network across samples.

**Ex vivo µCT analysis of long bones**

Ex vivo µCT imaging of long bones (humeri) was performed using a high-definition desktop µCT system (Bruker SkyScan 1272, Bruker microCT, Kontich, Belgium). Scans were performed using a tube voltage of 60 kVp and a tube current of 166 mA, with an exposure time of 1166 ms per projection. A 250 µm aluminum filter was applied to reduce beam-hardening effects. Images were acquired with an isotropic voxel size of 5 µm, corresponding to a voxel volume of $1.25 \times 10^{-7}$ mm³, and a field of view of 3 × 3.5 × 13 mm. Data acquisition was conducted in step-and-shoot mode over a 360° rotation, with a rotation step of 0.1°. Projections were acquired over a 360° rotation and reconstructed using NRecon software (Bruker). Quantitative morphometric analyses were conducted using Imalytics Preclinical (Gremse-IT GmbH, Aachen, Germany). ROIs were defined for quantitative analysis as follows: the proximal ROI was cylindrical, 3 mm in diameter and 3.5 mm in height, located at the proximal end; the diaphyseal ROI was cylindrical, 1 mm in diameter and 2 mm in height, centered on the mid-diaphysis to sample cortical bone; and the distal ROI was cylindrical, 3 mm in diameter and 3.5 mm in height, located at the distal end of the humerus. A qualitative overview of intramedullary bone buildup is provided in **Table S7**.

**Immunohistochemistry staining of paraffin-embedded tissue**

Humeri were embedded in paraffin following standard histological procedures. Formalin-fixed paraffin embedded (FFPE) blocks were cut into 5 µm thick slices. Tissue deparaffinization was performed at 60 °C for 2 h followed by xylol and ethanol serial dilution. Antigen retrieval was performed at 98 °C for 20 min with an antigen unmasking solution, citric acid based (H-3300-250 Vector Laboratories, Inc. Newark, United States) in PBS, followed by cooling in ice for 10 min. For antibody staining, endogenous peroxidase activity was blocked by using Bloxall endogenous blocking solution (Vector Laboratories, Inc. Newark, United States) for 1 h at RT, sections were incubated with the primary antibody (Origene, R1038) diluted in 12% BSA for 60 min at RM and overnight at 4°C. Subsequently, slides were washed three times with PBS for 5 min and incubated with secondary antibodies for 45 min. Thereafter, FFPE slides were washed with PBS and staining was developed with ImmPACT® DAB (Vector Laboratories, Inc. Newark, United States) according to the manufacturer's instructions. Hematoxylin was employed as counterstain for nuclei visualization, next slides were preserved using ethanol and xylol serial dilution and mounted with Vitro-Clud® (R. Langenbrinck GmbH, Emmendingen, Germany).

### Supplementary Tables

#### Table 1. List of all healthy donor and patient-derived samples.

| Sample ID | Diagnosis | Mutation | JAK2V617F VAF (%) | Additional mutations (%) | Treatment at sampling | Sample source | Material used | Experiment(s) |
|---|---|---|---|---|---|---|---|---|
| MPN-001 | PV | $JAK2^{V617F}$ | 53 | - | IFNa | PB | PBMC and $CD34^+$ | CFU, apoptosis assay, 3'-mRNASeq |
| MPN-002 | PV | $JAK2^{V617F}$ | 52 | ASXL1 21% 1 bp Dup (p.Ser892Phefs*2) | n.a. | PB | PBMC and $CD34^+$ | CFU, apoptosis assay |
| MPN-003 | PV | $JAK2^{V617F}$ | 78 | TET2 40% Q876* | n.a. | PB | PBMC | CFU, glucose uptake assay |
| MPN-004 | PV | $JAK2^{V617F}$ | 80 | - | n.a. | PB | PBMC | CFU, glucose uptake assay |
| MPN-005 | PV | $JAK2^{V617F}$ | 84 | - | Wait and watch | PB | PBMC | CFU, glucose uptake assay |
| MPN-006 | PV | $JAK2^{V617F}$ | 83 | - | Hydroxyurea | PB | PBMC and $CD34^+$ | CFU, apoptosis assay |
| MPN-007 | PV | $JAK2^{V617F}$ | 78 | TET2 39% 1 bp del (p.Phe57Serfs*10) | n.a. | PB | PBMC and $CD34^+$ | IF (mono and co-culture) |
| MPN-008 | PV | $JAK2^{V617F}$ | 51 | - | Ruxolitinib | PB | PBMC | IF (mono and co-culture) |
| MPN-009 | PV | $JAK2^{V617F}$ | 63 | - | Ruxolitinib | PB | PBMC | IF (mono and co-culture) |
| MPN-010 | PV | $JAK2^{V617F}$ | 71 | - | IFNa | PB | PBMC | IF (mono and co-culture) |
| MPN-011 | PV | $JAK2^{V617F}$ | 43 | TET2 5.3% Q235*, TET2 6.6% E1352K | Ruxolitinib | PB | PBMC | IF (mono and co-culture) |
| MPN-012 | PV | $JAK2^{V617F}$ | 57 | TET2 32% 2 bp del (p.His974Glnfs*8) | IFNa | PB | PBMC | IF (mono and co-culture) |
| MPN-013 | Post PV-MF | $JAK2^{V617F}$ | 63 | - | n.a. | PB | PBMC | IF (mono and co-culture) |
| MPN-014 | Post PV-MF | $JAK2^{V617F}$ | 78 | - | n.a. | PB | PBMC | IF (mono and co-culture) |
| MPN-015 | PV | $JAK2^{V617F}$ | 52 | - | Hydroxyurea | PB | $CD34^+$ | 3'-mRNAseq |
| MPN-016 | PV | $JAK2^{V617F}$ | 59 | ASXL1 30% E931* | n.a. | PB | $CD34^+$ | 3'-mRNAseq |
| MPN-017 | Post ET-MF | $JAK2^{V617F}$ | 59 | - | n.a. | PB | $CD34^+$ | 3'-mRNAseq |
| MPN-018 | Post PV-MF | $JAK2^{V617F}$ | 73 | - | Ruxolitinib | PB | $CD34^+$ | 3'-mRNAseq |
| MPN-019 | PMF | $JAK2^{V617F}$ | 45 | - | Wait and watch | PB | $CD34^+$ | 3'-mRNAseq |
| HD-001 | - | - | - | - | - | PB | PBMC | CFU |
| HD-002 | - | - | - | - | - | PB | PBMC | CFU |
| HD-003 | - | - | - | - | - | PB | PBMC | CFU |
| HD-004 | - | - | - | - | - | Leukotrap | $CD34^+$ | Apoptosis assay |
| HD-005 | - | - | - | - | - | Leukotrap | $CD34^+$ | Apoptosis assay |
| HD-006 | - | - | - | - | - | Leukotrap | $CD34^+$ | Apoptosis assay |
| HD-004 | - | - | - | - | - | Leukotrap | $CD34^+$ | 3'-mRNAseq |
| HD-005 | - | - | - | - | - | Leukotrap | $CD34^+$ | 3'-mRNAseq |
| HD-006 | - | - | - | - | - | Leukotrap | $CD34^+$ | 3'-mRNAseq |

**Table 2. List of primers.**

| Primer ID | Target gene | Species | Application | Forward primer sequence (5′–3′) | Reverse primer sequence (5′–3′) |
|---|---|---|---|---|---|
| 001 | *ACTA2* | Human | qPCR | TCCTTCATCGGGATGGAGTCT | TACATAGTGGTGCCCCCTGA |
| 002 | *FN1* | Human | qPCR | TCGCAGCTTCGAGATCAGTG | CCCCTCTTCATGACGCTTGT |
| 003 | *GLI1* | Human | qPCR | GAGCCAGAAGTTGGGACCTC | CCTCGCTCCATAAGGCTCAG |
| 004 | *MT-ATP6* | Human | qPCR | CGTACGCCTAACCGCTAACA | AGGCGACAGCGATTTCTAGG |
| 005 | *IL1B* | Murine | qPCR | AGATGGTCAATGGCAGAACTGT | TGATGTGCTGCTGCGAGATT |
| 006 | *IL1RL1* | Murine | qPCR | GGATTGAGGTTGCTCTGTTCTGG | TCGGGCAGAGTGTGGTGAACAA |
| 007 | *GAPDH* | Murine | qPCR | CCAATACGGCCAAATCCG | TTGTGCAGTGCCAGCCTC |

**Table 3. List of flow cytometry antibodies.**

| ID | Antibody | Company | Cat no. | Reactivity | Dilution |
|---|---|---|---|---|---|
| 001 | CD45.1-PB | Biolegend | 110722 | Anti-mouse | 1:200 |
| 002 | CD45.2-PE | Biolegend | 109808 | Anti-mouse | 1:200 |
| 003 | CD45.2-FITC | Biolegend | 109806 | Anti-mouse | 1:200 |
| 004 | CD3-PB | Biolegend | 100214 | Anti-mouse | 1:100 |
| 005 | Ter119-APC | Biolegend | 116212 | Anti-mouse | 1:100 |
| 006 | CD71-PE | Biolegend | 113808 | Anti-mouse | 1:100 |
| 007 | CD3-PE-Cy5 | BD Biosciences | 555276 | Anti-mouse | 1:200 |
| 008 | CD4-PE-Cy5 | BD Biosciences | 553050 | Anti-mouse | 1:200 |
| 009 | CD8a-PE-Cy5 | eBiosciences | 15-0081-82 | Anti-mouse | 1:200 |
| 010 | B220-PE-Cy5 | Biolegend | 103210 | Anti-mouse | 1:200 |
| 011 | Gr1-PE-Cy5 | Biolegend | 108410 | Anti-mouse | 1:200 |
| 012 | CD11b-PE-Cy5 | Biolegend | 101210 | Anti-mouse | 1:200 |
| 013 | Ter119-PE-Cy5 | Biolegend | 116210 | Anti-mouse | 1:200 |
| 014 | CD48-PB | Biolegend | 103418 | Anti-mouse | 1:100 |
| 015 | Sca1-PE-Cy7 | Biolegend | 122514 | Anti-mouse | 1:100 |
| 016 | CD150-APC | Biolegend | 115910 | Anti-mouse | 1:100 |
| 017 | CD117-APC-Cy7 | Biolegend | 105826 | Anti-mouse | 1:100 |
| 018 | CD34-eFluor™ 450 | ThermoScientific | 48-0341-82 | Anti-mouse | 1:50 |
| 019 | CD16/32-APC | Biolegend | 156608 | Anti-mouse | 1:100 |

**Table 4. List of primary antibodies used for mono and co-culture.**

| ID | Antibody | Reactivity | Company | Cat no. | Dilution |
|---|---|---|---|---|---|
| 001 | alpha-SMA | Anti-Mouse | R&D Systems | MAB1420 | 1:500 |
| 002 | phospho-SMAD2 (Ser465/467)/SMAD3 (Ser423/425) | Anti-Rabbit | Cell Signaling | 8828 | 1:200 |

**Table 5. List of secondary antibodies used for mono and co-culture.**

| ID | Antibody | Conjugate | Reactivity | Company | Cat no. | Dilution |
|---|---|---|---|---|---|---|
| 001 | Goat anti-mouse IgG (H+L) | Alexa Fluor™ Plus 488 | Mouse | Thermo Fisher | A32723 | 1:100 |
| 002 | Goat anti-rabbit IgG (H+L) | Alexa Fluor™ 750 | Rabbit | Thermo Fisher | A21039 | 1:100 |

**Table 6. Progenitor medium composition.**

| ID | Component | Final Conc. | Cat no. | Company |
|---|---|---|---|---|
| 001 | IMDM | 50% | 12440-053 | Gibco |
| 002 | Ham's F-12 nutrient mix | 50% | 11765-054 | Gibco |
| 003 | Human albumin (Albutein 200 g/L) | 0.5% | 17910338 | Grifols |
| 004 | GlutaMAX™ | 2 mM | 15140-130 | Gibco |
| 005 | Chem. Defined Lipids | 2 mM | 11905-031 | Gibco |
| 006 | L-ascorbic acid | 50 µg/mL | A92902 | Sigma-Aldrich |
| 007 | Holo-transferrin | 6 µg/mL | T0665 | Sigma-Aldrich |
| 008 | 1-thioglycerol | 400 µM | M1753 | Sigma-Aldrich |

**Table 7. Summary of animal experiments with observed sclerosis.**

| Group | Mouse | Sex | Sclerosis | Prox | Diaphysis | Distal | Notes |
|---|---|---|---|---|---|---|---|
| **$JAK2^{WT}$ vehicle** | 1 | m | | | | | |
| | 2 | m | | | | | |
| | 3 | m | X | X | | X | fine mesh, init. |
| | 4 | f | | | | | |
| | 5 | f | | | | | |
| | 6 | f | | | | | |
| **$JAK2^{V617F}$ vehicle** | 7 | f | X | | X | X | central canal |
| | 8 | m | X | | | X | spots |
| | 9 | m | X | | X | X | central canal |
| | 10 | m | X | X | | | |
| | 11 | m | X | | | X | one spot |
| | 12 | m | | | | | |
| | 13 | f | X | | X | X | spots |
| | 14 | f | X | X | X | X | diffused, spots |
| | 15 | f | X | | X | | one spot |
| **Melatonin** | 16 | f | X | | X | X | central canal |
| | 17 | f | X | | X | X | central canal |
| | 18 | f | X | X | X | X | diffused |
| | 19 | f | | | | | |
| | 20 | f | X | | X | | one spot |
| | 21 | m | X | | | X | endosteal react. |
| | 22 | m | X | | X | X | central canal |
| | 23 | m | | | | | |
| **Ruxolitinib** | 24 | m | X | | | X | |
| | 25 | m | | | | | |
| | 26 | m | X | | X | X | spots |
| | 27 | m | X | | X | X | spots |

| | | | | | | | |
|---|---|---|---|---|---|---|---|
| | 28 | m | X | | | X | |
| | 29 | f | X | X | X | X | |
| **MT+Ruxolitinib** | 30 | f | X | | X | | central canal |
| | 31 | m | | | | | |
| | 32 | m | X | | X | X | central canal |
| | 33 | m | X | | X | X | |
| | 34 | m | X | | | X | one spot |
| | 35 | m | X | | X | X | |
| **Nano-MT** | 36 | m | | | | | |
| | 37 | m | X | | X | X | spots, central canal |
| | 38 | m | X | X | X | | fine mesh, init. |
| | 39 | m | X | | X | X | central canal |
| | 40 | f | | | | | |
| | 41 | f | X | | X | | central canal |
| | 42 | f | X | | X | | central canal |
| | 43 | f | X | | X | X | spots, central canal |

### Supplementary Figures

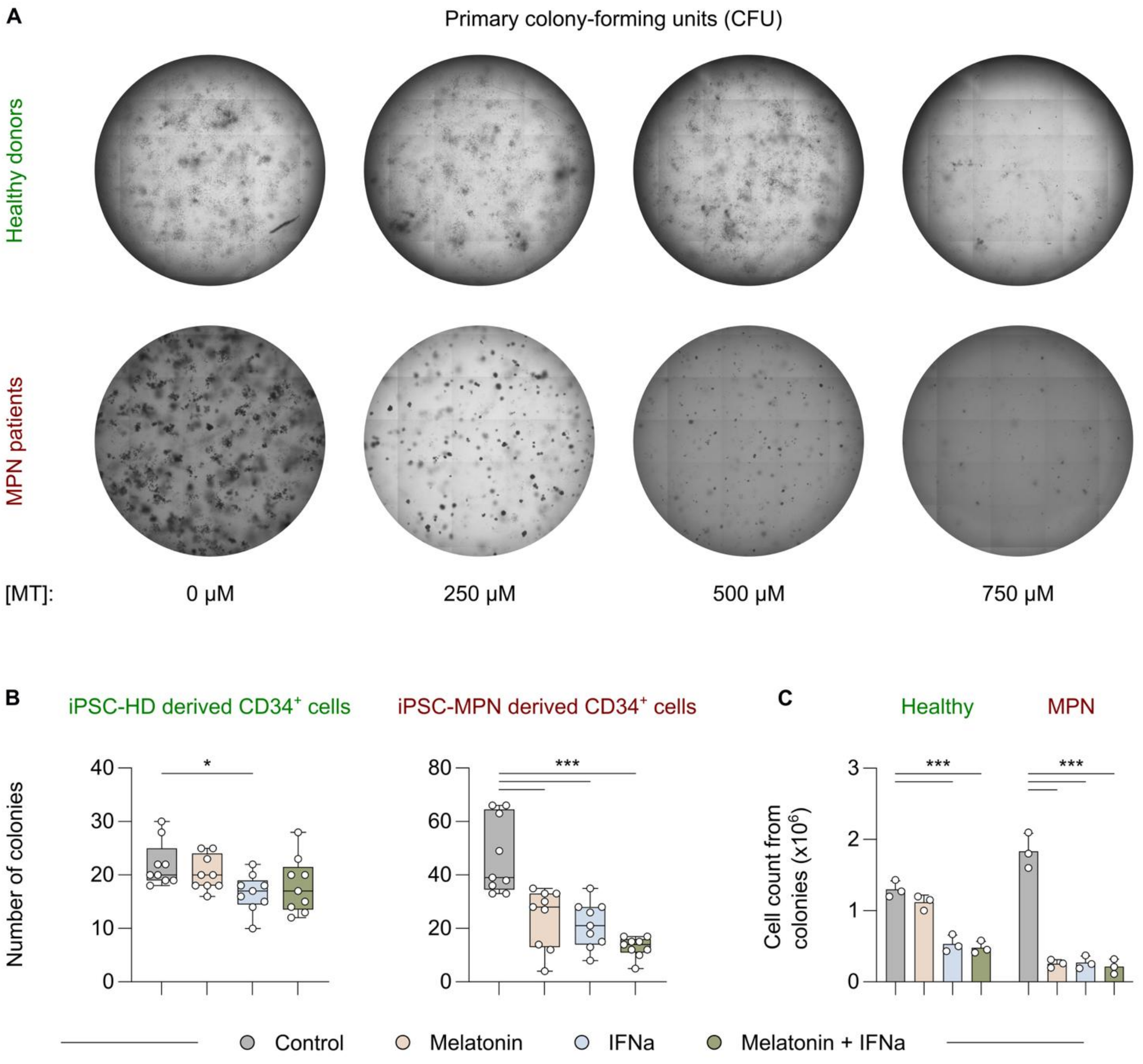


**Figure S1. MT reduces clonogenic growth of MPN PBMCs and $CD34^+$ cells in vitro. (A)** Representative images of total colonies from PBMC-derived CFU assays of healthy donor and MPN patients cultured with increasing concentrations of melatonin (0, 250, 500, 750 μM). The images are acquired at 10x magnification. **(B)** Total colony numbers from $CD34^+$ enriched HD (left) and CALR del52het01 MPN patients (right) after 14 days of CFU assay treated with Vehicle control (DMSO, 0.25%), Melatonin (500 μM), IFN-α (1000 U/mL), and combination (Melatonin + IFN-α). Kruskal-Wallis test with Dunn's corrections for multiple comparisons was performed. **(C)** Total viable cell counts from (B); Two-Way ANOVA with Šídák's test for multiple comparisons was used after passing the Shapiro-Wilk normality test; $^*P < 0.05$, $^{***}P < 0.001$.

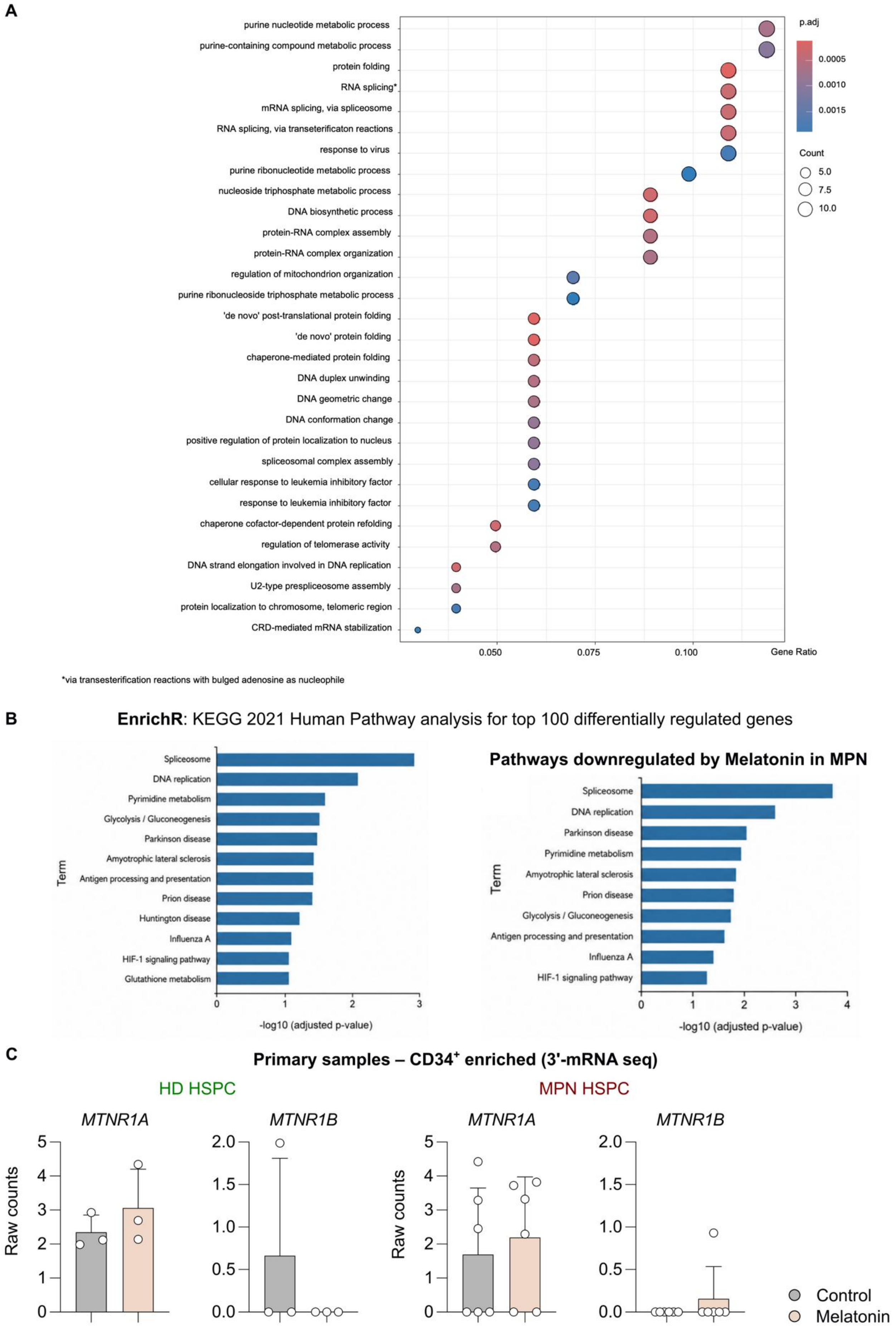
A
purine nucleotide metabolic process
purine-containing compound metabolic process
protein folding
RNA splicing*
mRNA splicing, via spliceosome
RNA splicing, via transeterificaton reactions
response to virus
purine ribonucleotide metabolic process
nucleoside triphosphate metabolic process
DNA biosynthetic process
protein-RNA complex assembly
protein-RNA complex organization
regulation of mitochondrion organization
purine ribonucleoside triphosphate metabolic process
'de novo' post-translational protein folding
'de novo' protein folding
chaperone-mediated protein folding
DNA duplex unwinding
DNA geometric change
DNA conformation change
positive regulation of protein localization to nucleus
spliceosomal complex assembly
cellular response to leukemia inhibitory factor
response to leukemia inhibitory factor
chaperone cofactor-dependent protein refolding
regulation of telomerase activity
DNA strand elongation involved in DNA replication
U2-type prespliceosome assembly
protein localization to chromosome, telomeric region
CRD-mediated mRNA stabilization
0.050
0.075
0.100
Gene Ratio
p.adj
0.0005
0.0010
0.0015
Count
5.0
7.5
10.0
*via transesterification reactions with bulged adenosine as nucleophile
B
EnrichR: KEGG 2021 Human Pathway analysis for top 100 differentially regulated genes
Spliceosome
DNA replication
Pyrimidine metabolism
Glycolysis / Gluconeogenesis
Parkinson disease
Amyotrophic lateral sclerosis
Antigen processing and presentation
Prion disease
Huntington disease
Influenza A
HIF-1 signaling pathway
Glutathione metabolism
Term
-log10 (adjusted p-value)
Pathways downregulated by Melatonin in MPN
Spliceosome
DNA replication
Parkinson disease
Pyrimidine metabolism
Amyotrophic lateral sclerosis
Prion disease
Glycolysis / Gluconeogenesis
Antigen processing and presentation
Influenza A
HIF-1 signaling pathway
Term
-log10 (adjusted p-value)
C
Primary samples – CD34+ enriched (3'-mRNA seq)
HD HSPC
MPN HSPC
MTNR1A
MTNR1B
MTNR1A
MTNR1B
Raw counts
Raw counts
Control
Melatonin

**Figure S2. MT reduces key metabolic pathways in MPN independent of the MT receptor activation *in vitro*. (A)** Dot plot summarizing pathway-level changes derived from 3′ mRNA-seq of $CD34^+$ hematopoietic stem and progenitor cells (HSPC) treated with melatonin versus vehicle control (0.25% DMSO). Each dot represents one pathway; dot size indicates the number of genes contributing to the enrichment signal and dot color indicates significance (adj. P-value). Pathway enrichment was computed from differentially expressed genes. **(B)** KEGG pathway enrichment analysis of the same 3′ mRNA-seq differential expression dataset (melatonin vs vehicle in treated MPN samples) performed using EnrichR. The plot displays significantly enriched KEGG pathways ranked by $-\log_{10}$FC (adj. P-value). **(C)** Normalized read counts for the melatonin receptor genes *MTNR1A* and *MTNR1B* obtained from the same 3′ mRNA-seq dataset ($CD34^+$ HSPC), n = 3 (HD), n = 6 (MPN). Data are presented as Mean of Raw Counts ± SD.

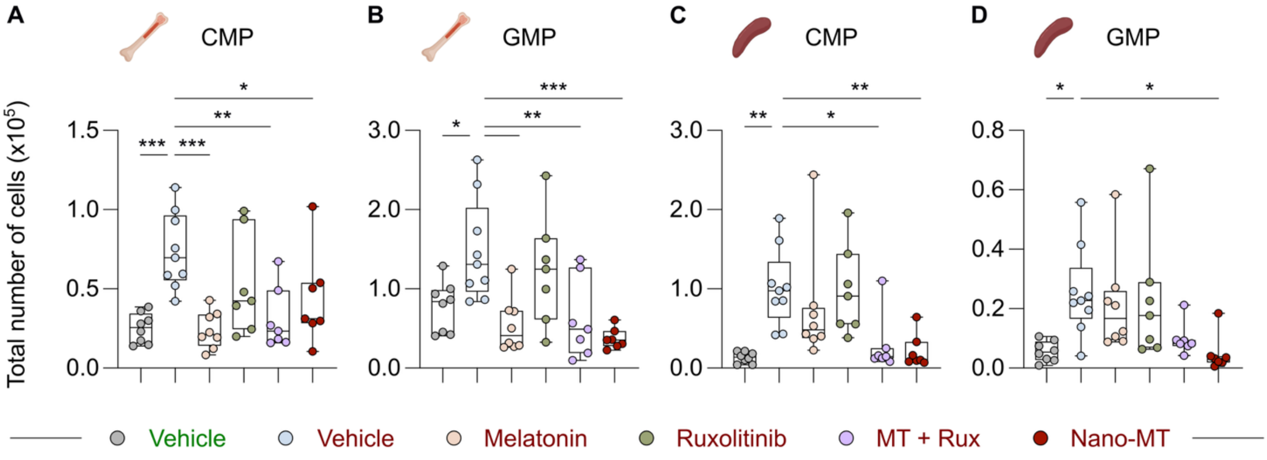


**Figure S3. Nano-MT reduces mutant myeloid progenitor output in bone marrow and spleen. (A,B)** Absolute numbers of myeloid progenitor populations in BM, including common myeloid progenitors (CMP) and granulocyte-monocyte progenitors (GMP) across $JAK2^{WT}$ treated with vehicle, and $JAK2^{V617F}$ treated with vehicle, melatonin (MT), ruxolitinib (Rux), melatonin + ruxolitinib combination, and liposomal melatonin (Nano-MT). **(C,D)** Absolute numbers of splenic CMP and GMP with the same treatment groups are shown. Statistical analysis was conducted via One-Way ANOVA with Dunnett's test for multiple comparisons; *P < 0.05, **P < 0.01, ***P < 0.001.

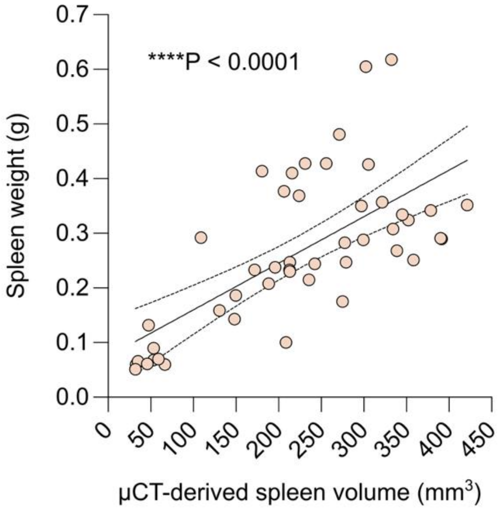


**Figure S4. Correlation between µCT-derived spleen size and terminal spleen weight.** Scatter plot showing the correlation between spleen volume measured by *in vivo* µCT and *ex vivo* spleen weight measured at sacrifice time point. The solid line indicates the simple linear regression fit, and dotted lines indicate the 95% confidence interval. The correlation between µCT-derived spleen volume and spleen weight was assessed via Pearson correlation; ****P < 0.0001, r = 0.684, $R^2$ = 0.468, n = 47.

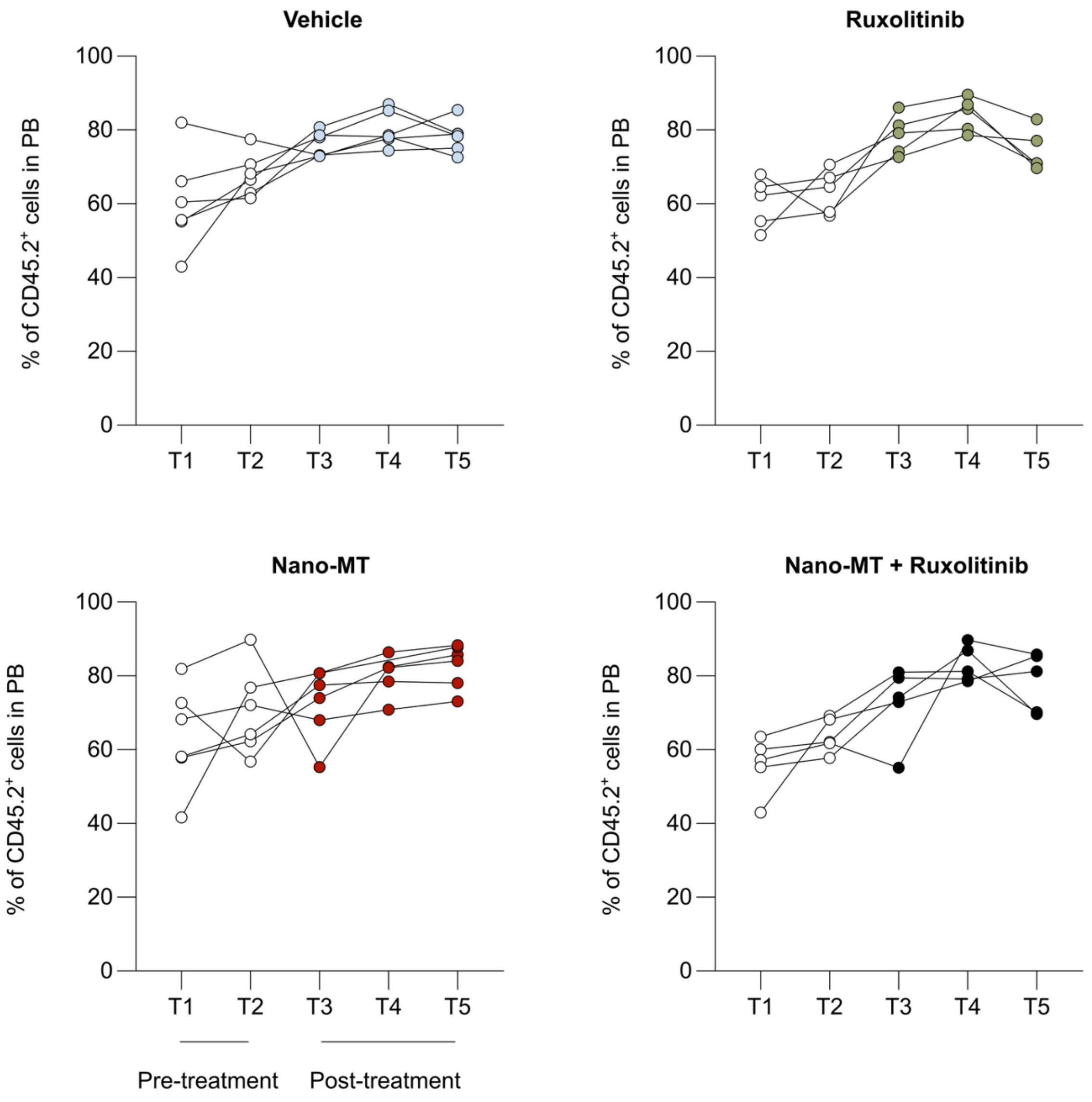


**Figure S5. Longitudinal peripheral blood CD45.2$^+$ engraftment during treatment.** Percentage of CD45.2$^+$ cells in peripheral blood (PB) over time in JAK2$^{V617F}$ mice treated with vehicle, ruxolitinib, liposomal melatonin (Nano-MT), and Nano-MT + ruxolitinib combination. T1 and T2 represent pre-treatment time points, whereas T3, T4, and T5 correspond to 2, 4, and 6 weeks of treatment, respectively. Individual mice values are connected by lines. White circles indicate pre-treatment measurements, and colored circles indicate measurements upon treatment initiation.

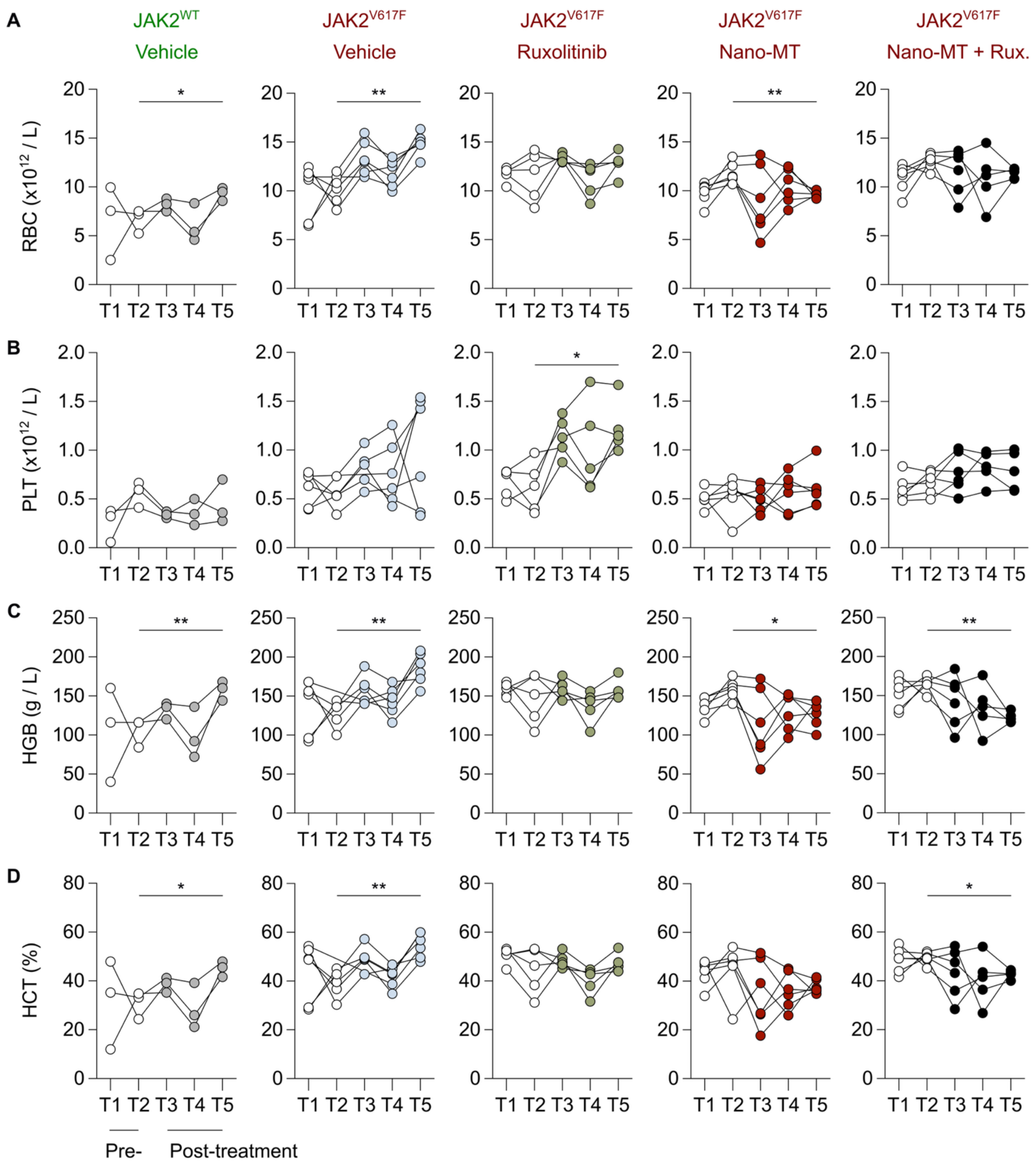


**Figure S6. Longitudinal changes in peripheral blood parameters during treatment. (A-D)** Longitudinal peripheral blood measurements of red blood cell count (RBC), platelets (PLT), hemoglobin (HGB), hematocrit (HCT) in JAK2$^{WT}$ mice treated with vehicle, and JAK2$^{V617F}$ mice treated with vehicle, ruxolitinib (Rux), liposomal melatonin (Nano-MT), and Nano-MT + ruxolitinib combination. T1 and T2 represent pre-treatment time points, whereas T3, T4, and T5 correspond to 2, 4, and 6 weeks of treatment, respectively. Individual mouse values are connected by lines. White circles indicate pre-treatment measurements, and colored circles indicate measurements upon treatment initiation. Statistical comparisons were performed between T2 and T5 within each group via a paired t-test; * $P < 0.05$; ** $P < 0.01$.
